# Life without cytoplasm: sperm sustain protein production utilizing cytoplasmic droplets and mitochondria as translational apparatuses

**DOI:** 10.64898/2026.08.10.744062

**Authors:** Zhuqing Wang, Hetan Wang, Rubens Daniel Miserani Magalhães, Sheng Chen, Siying Meng, Dayton Morris, Sarah Crane, Hayden McSwiggin, Anne Elizabeth Yan, Saayli Khambekar, Brian Nguyen, Huili Zheng, Wei Yan

**Author notes:** These authors contributed equally (Zhuqing Wang, Hetan Wang).

## Abstract

Sperm present a unique paradox: heavily compacted chromatin silences transcription, and loss of cytoplasm during spermiation eliminates the conventional translational apparatus, yet sperm require ∼10 days of protein-demanding epididymal maturation to acquire motility and fertilization competence. How sperm produce the necessary proteins has remained enigmatic. Here, we show that mammalian sperm sustain protein synthesis through a biphasic translational program. In testicular sperm, a residual cytoplasmic droplet retains the full complement of translational machinery, including ribosomes and intact tRNAs, and supports nascent protein synthesis. As sperm enter the epididymis, this droplet is progressively fragmented, and protein synthesis shifts to the midpiece, where mitochondrial translation machinery becomes active. Proteomic and functional analyses indicate that both systems contribute to sperm maturation. Inhibiting cytoplasmic translation in testicular sperm or mitochondrial translation in epididymal sperm abolishes motility. These findings reveal how cytoplasm-free sperm overcome transcriptional silence to complete maturation and support male fertility.

**Graphic Abstract:** 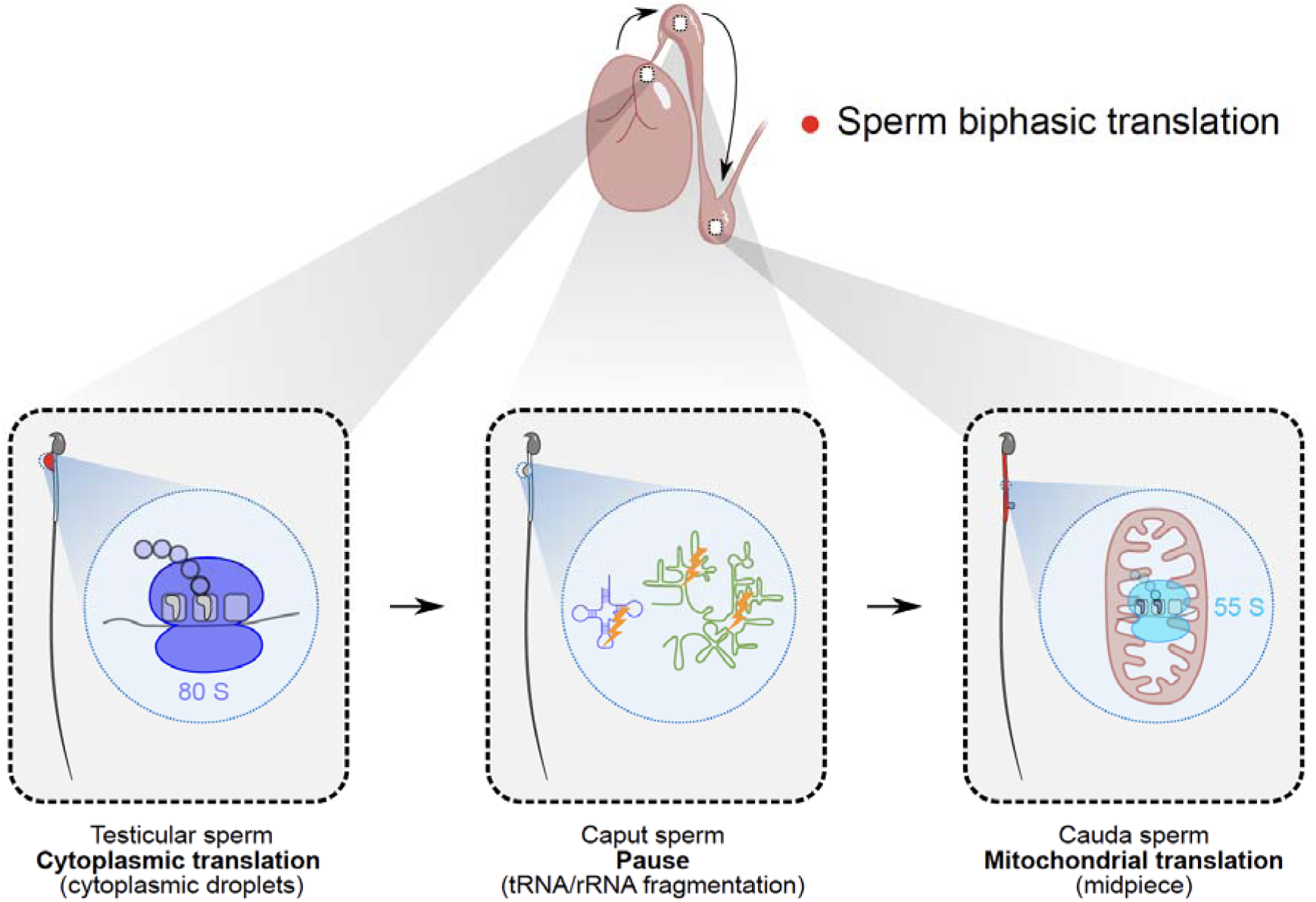

## Introduction

Spermiogenesis, the terminal phase of spermatogenesis, transforms round spermatids into morphologically mature elongated spermatids through a sequence of dramatic cellular remodeling events, including nuclear condensation, acrosome biogenesis, and flagellum assembly^1^. Central to this transformation is the wholesale replacement of histones with protamine, producing highly compacted chromatin that renders the sperm genome transcriptionally inert^2,3^. The associated loss of most cytoplasm, extruded as residual bodies phagocytosed by Sertoli cells during spermiation, has long led to the assumption that spermatozoa are also translationally silent^4,5^. This dogma has persisted despite compelling evidence that sperm must undergo extensive further maturation in the epididymis to acquire fertilization competence. During ∼10 days of epididymal transit, sperm undergo substantial structural and biochemical transformation: plasma membrane remodeling, chromatin stabilization via disulfide cross-linking, acquisition of progressive motility, and extensive reshaping of their small RNA cargo^6–10^. These processes demand specific proteins, which must originate either from preloaded reserves or from *de novo* synthesis. Yet the source and mechanistic basis of protein production in maturing sperm have remained poorly understood, representing a fundamental gap in reproductive biology.

A conspicuous feature of testicular and epididymal spermatozoa that has long resisted functional characterization is the cytoplasmic droplet (CD), a remnant of cytoplasm first described by Retzius in 1909^11^. CDs are initially positioned at the sperm neck and migrate distally along the flagellum as sperm progress through the epididymis^12^. In mice, more than 80% of caput sperm retain CDs, whereas only 40–50% of cauda sperm do, reflecting gradual droplet loss during maturation^12^. Persistently retained CDs in ejaculated sperm are a recognized marker of defective maturation^12,13^. Proteomic analyses have shown that CDs are enriched in proteins involved in energy metabolism, phosphorylation signaling, cytoskeletal organization, membrane trafficking, and stress response^9,12,14^. Despite this suggestive composition, CD function has remained enigmatic. Genetic studies linking CD malformation to infertility, as in the loss of *Spem1*^15^, *Sypl1*^16^, *Arrdc5*^17^, or *Tex38*^17,18^, underscore their physiological importance, yet the mechanistic basis of CD function has not been established. Several questions remain unresolved: why do early spermatozoa harbor CDs that are progressively shed during maturation, and why do sperm tRNAs and rRNAs undergo fragmentation during epididymal transit?

Several lines of indirect evidence have pointed toward a translational function for CDs. First, ultrastructural analyses have revealed that CDs are enriched in monosomes, polysomes, and the Golgi apparatus^19–21^, suggesting active translation and protein folding. Second, CDs contain abundant mRNAs and circular RNAs with coding potential^22^, as well as a dynamic complement of small RNAs, including tRNA-derived small RNAs (tsRNAs) and rRNA-derived small RNAs (rsRNAs)^10,23–26^. Our previous work demonstrated that tsRNAs and rsRNAs are actively trafficked between CDs and sperm during epididymal transit^10^, and several reports have established their roles in translational regulation^27–32^. Third, sperm contain large, intact mRNAs^22,33–35^ that represent potential translational substrates. Together, these observations suggested that CDs might function as a translational apparatus, utilizing preloaded RNAs to synthesize proteins *de novo* to meet the requirements for maturation.

We therefore set out to test the hypothesis that CDs function as a transient translational organelle during epididymal sperm maturation. Here, we demonstrate that sperm employ a striking biphasic translational strategy: testicular sperm exploit a complete cytoplasmic translational machinery housed within CDs, which is then silenced as sperm enter the caput epididymis coincident with tRNA and rRNA fragmentation, and subsequently replaced by mitochondrial translation in cauda sperm. Using SUnSET, Ribo-lite, and BONCAT approaches, we map the translatome and nascent proteome at each stage. Critically, we demonstrate that pharmacological blockade of cytoplasmic translation by intra-rete testis injection of cycloheximide almost completely abolishes sperm motility and that pre-treatment of cauda sperm with chloramphenicol prior to HTF-induced activation abolishes progressive motility, establishing direct causal requirements for both phases of the biphasic translational program in sperm functional competence. These findings redefine CDs as essential biosynthetic organelles and reveal an unanticipated mechanism by which transcriptionally silent, cytoplasm-free sperm maintain their translational capacity during maturation and possibly into fertilization.

## Results

### Protein composition undergoes dynamic remodeling as sperm transit the epididymis

If CDs function to synthesize novel proteins during sperm maturation, protein composition should change dynamically as sperm progress from the testis through the epididymis. To test this, we first characterized protein profiles in testicular, caput, and cauda epididymal sperm by LC-MS/MS-based proteomics. To ensure high-purity sperm preparations free of somatic cell contamination, we employed the Percoll centrifugation protocol combined with somatic cell lysis buffer (SCLB) described previously^10^, achieving purities of 98.5% (testicular), 98.7% (caput), and 99.6% (cauda) sperm.

Because conventional RIPA lysis buffer fails to efficiently lyse sperm heads due to extensive disulfide bonds, we optimized a lysis protocol that incorporated additional SDS and 2-mercaptoethanol, followed by sonication and heating (Figure S1). Proteomic analyses identified 3,597 proteins in testicular sperm, 2,750 in caput sperm, and 2,391 in cauda sperm (Table S1). Principal component analysis demonstrated clear separation of all three populations, with caput and cauda sperm clustering more closely together than with testicular sperm (Figure 1A), indicating progressive remodeling during epididymal transit. Pairwise comparisons identified 445 upregulated and 672 downregulated proteins between testicular and caput sperm (Figure 1B; Table S1; Log_2_ fold change ≥ 2, p < 0.022), and 95 upregulated and 109 downregulated proteins between caput and cauda sperm (Figure 1C; Table S1; Log_2_ fold change ≥ 2, p < 0.0054). During the testis-to-caput transition, upregulated proteins were predominantly involved in carboxylic acid metabolism, tricarboxylic acid (TCA) cycle, and lipid metabolism, consistent with increasing energetic demands, while downregulated proteins were enriched for functions in translation, spermatogenesis, and sperm motility (Figures 1B and 1D; Table S2). During caput-to-cauda transit, energy metabolism proteins continued to rise, while RNA metabolic process proteins declined (Figures 1C and 1D; Table S2). Strikingly, 60 proteins were upregulated from testis to caput but subsequently downregulated from caput to cauda (Figures 1B-1D; Table S1), indicating a cohort of proteins specifically required during the caput phase; GO enrichment analysis showed these are predominantly RNA metabolism proteins (Figures 1B-1D). These dynamics are consistent with our prior observation that caput sperm exhibit a more diverse RNA profile than testicular or cauda sperm^10^.

**Figure 1.**
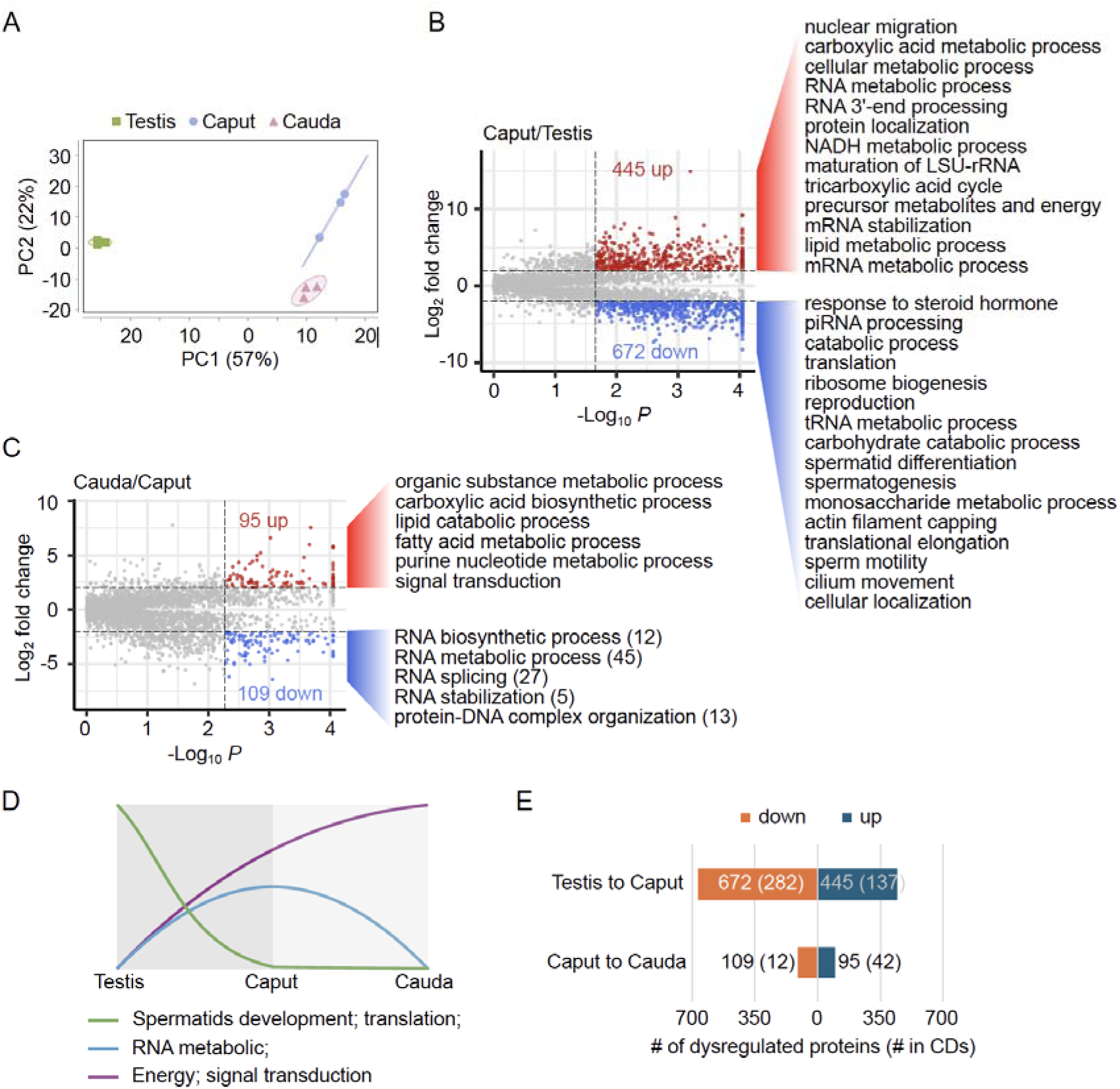
Proteins undergo dynamic changes as sperm transit from the testis to the epididymis. (A) Principal component analysis (PCA) of testicular, caput, and cauda sperm proteomics. (B) Volcano plot and GO term enrichment of dysregulated proteins between testicular and caput sperm. Upregulated proteins (red) are enriched for RNA metabolic processes and energy metabolism; downregulated proteins (blue) are enriched for translation, spermatogenesis, and sperm motility. (C) Volcano plot and GO term enrichment of dysregulated proteins between caput and cauda sperm. (D) Schematic summarizing the major GO terms enriched in testicular, caput, and cauda sperm proteomics. (E) Bar graph showing the number of dysregulated proteins during sperm epididymal transit (left bar) and the overlap with proteins identified in CD proteomics (numbers in parentheses).

Given that CDs are enriched in energy-metabolism proteins^14^, we hypothesized that CDs represent both the source of newly acquired proteins and the destination for proteins being shed. Comparing the 1,833 CD-resident proteins identified previously (Table S3) with the dysregulated proteins during epididymal transit, we found that >45% of all dysregulated proteins are present in CD proteomics (Figure 1E). This substantial overlap strongly suggests that CDs mediate the dynamic protein remodeling accompanying sperm epididymal maturation.

### A biphasic translational program switches from cytoplasmic to mitochondrial translation during epididymal transit

To determine whether dynamically changing proteins are preloaded or newly synthesized, we employed the Surface Sensing of Translation (SUnSET) approach^36^, which uses puromycin, a structural analog of tyrosyl-tRNA (Figure 2A), that is incorporated into elongating polypeptides and thus marks sites of active translation (Figure 2B)^37^. Following puromycin treatment of purified testicular, caput, and cauda sperm, immunofluorescence (IF) with anti-puromycin and anti-LDHC (a CD and sperm flagella marker^10,14^) revealed that translational activity in testicular sperm is concentrated at CDs (Figure 2C). Signal intensity was strongest in elongated spermatids and early testicular sperm and declined progressively during testicular transit. Puromycin signals were absent in caput sperm but reappeared in cauda sperm, where they localized specifically to the midpiece, the site of mitochondrial concentration, consistent with prior evidence of mitochondrial translational activity in mature sperm^5^.

**Figure 2.**
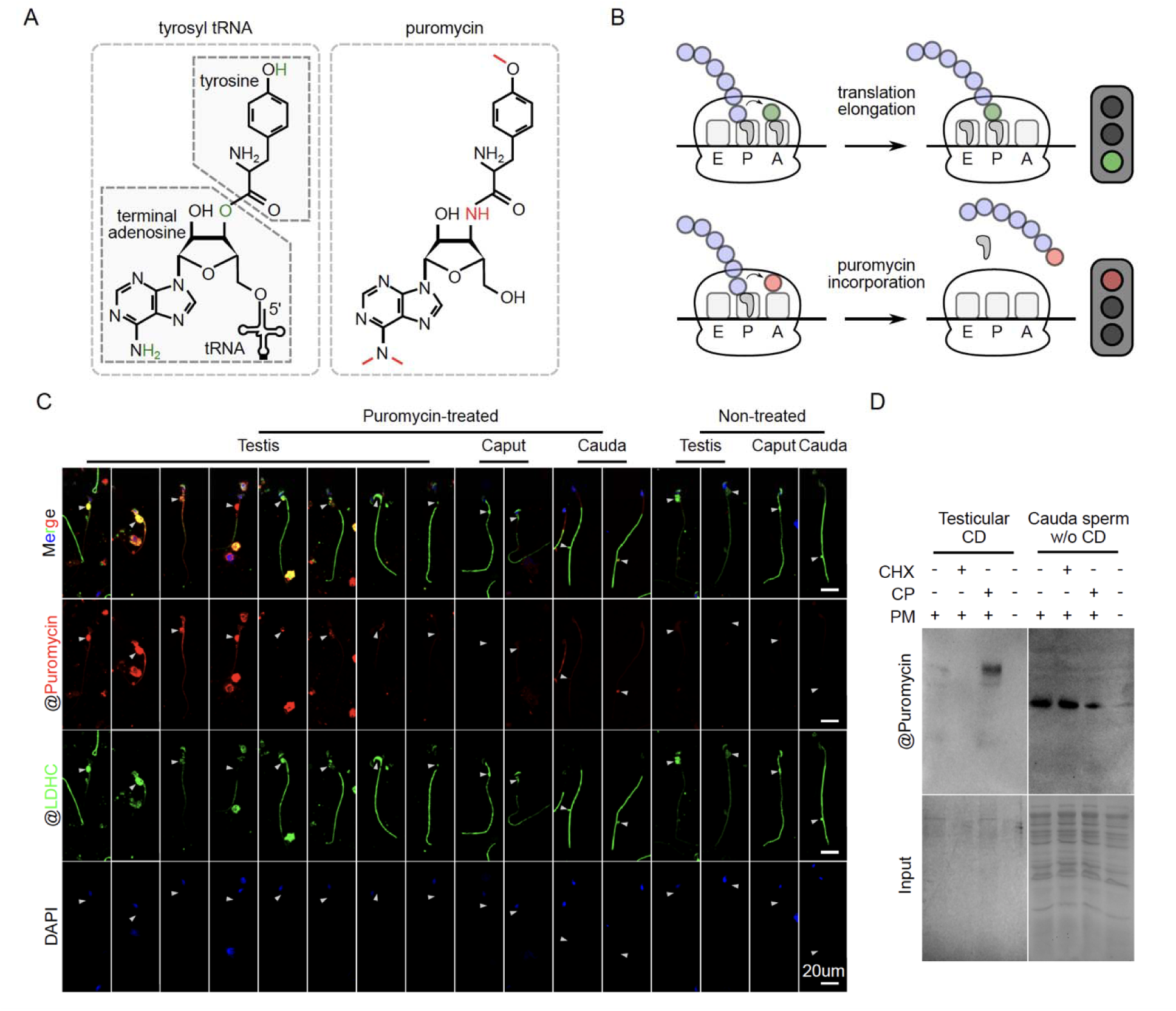
Sperm translate proteins in testicular CDs, cease translation in the caput epididymis, and resume translation via mitochondria in cauda sperm. (A) Chemical structures of tyrosyl-tRNA and puromycin, with structural differences highlighted in green and red, respectively. (B) Schematic of translation elongation and puromycin incorporation. P-site, peptidyl; A-site, aminoacyl; E-site, exit. Polypeptide, tyrosyl-tRNA, and puromycin are indicated in purple, green, and red, respectively. (C) Immunofluorescence of puromycin incorporation in testicular, caput, and cauda sperm. Puromycin (red), LDHC (green), DAPI (blue). Scale bar = 20 μm. (D) Western blot of puromycin incorporation in testicular CDs and CD-less cauda sperm treated with cycloheximide (CHX), chloramphenicol (CP), or no treatment (NT).

Because puromycin labeling cannot precisely report subcellular translation sites^38^, we purified testicular, caput, and cauda CDs and CD-less sperm separately and performed puromycin incorporation assays in EKRB buffer (Figure S2). Strong puromycin incorporation signals were detected in purified testicular CDs, but not in caput or cauda CDs (Figures S3A and S3B), confirming that testicular CDs alone possess *de novo* cytoplasmic translational activity. CD-less testicular and caput sperm showed weak incorporation, while CD-less cauda sperm showed strong incorporation confined to a discrete band at approximately 60 kDa, suggesting production of specific proteins (Figure S3). To distinguish cytoplasmic from mitochondrial translation, we used selective pharmacological inhibitors. Pretreatment of testicular CDs with cycloheximide (CHX, a cytoplasmic translation inhibitor) nearly abolished puromycin incorporation, while chloramphenicol (CP, a mitochondrial translation inhibitor) had no effect (Figure 2D). Conversely, in CD-less cauda sperm, CHX had no effect on translation, whereas CP significantly reduced it (Figure 2D). These results establish that testicular sperm use cytoplasmic translational machinery in CDs, while cauda sperm switch to mitochondrial translation.

### Testicular CDs, but not caput or cauda CDs, contain intact cytoplasmic translational machinery

Having established that testicular CDs are the principal site of cytoplasmic translation in maturing sperm, we sought to characterize the molecular components of this machinery. CD proteomics identified 117 proteins related to cytoplasmic translation among the 1,833 total CD-resident proteins, spanning six functional categories: 31 (26%) 60S ribosomal proteins, 26 (22%) translation initiation factors, 23 (20%) 40S ribosomal proteins, 20 (17%) tRNA ligases, 8 (7%) elongation factors, and 9 (8%) others (Figure 3A; Table S3). Western blot validation of seven representative proteins across these categories, RPL12, RPLP0, EIF3A, RPS3, WARS, EEF2, and RRBP1, confirmed their expression in whole testis and testicular CDs, but near-absence in SCLB-treated testicular sperm (which lack CDs) and in caput and cauda sperm (Figure 3B). Immunofluorescence corroborated strong CD-specific localization of EIF3A and RPL12 in testicular sperm (Figure 3C).

**Figure 3.**
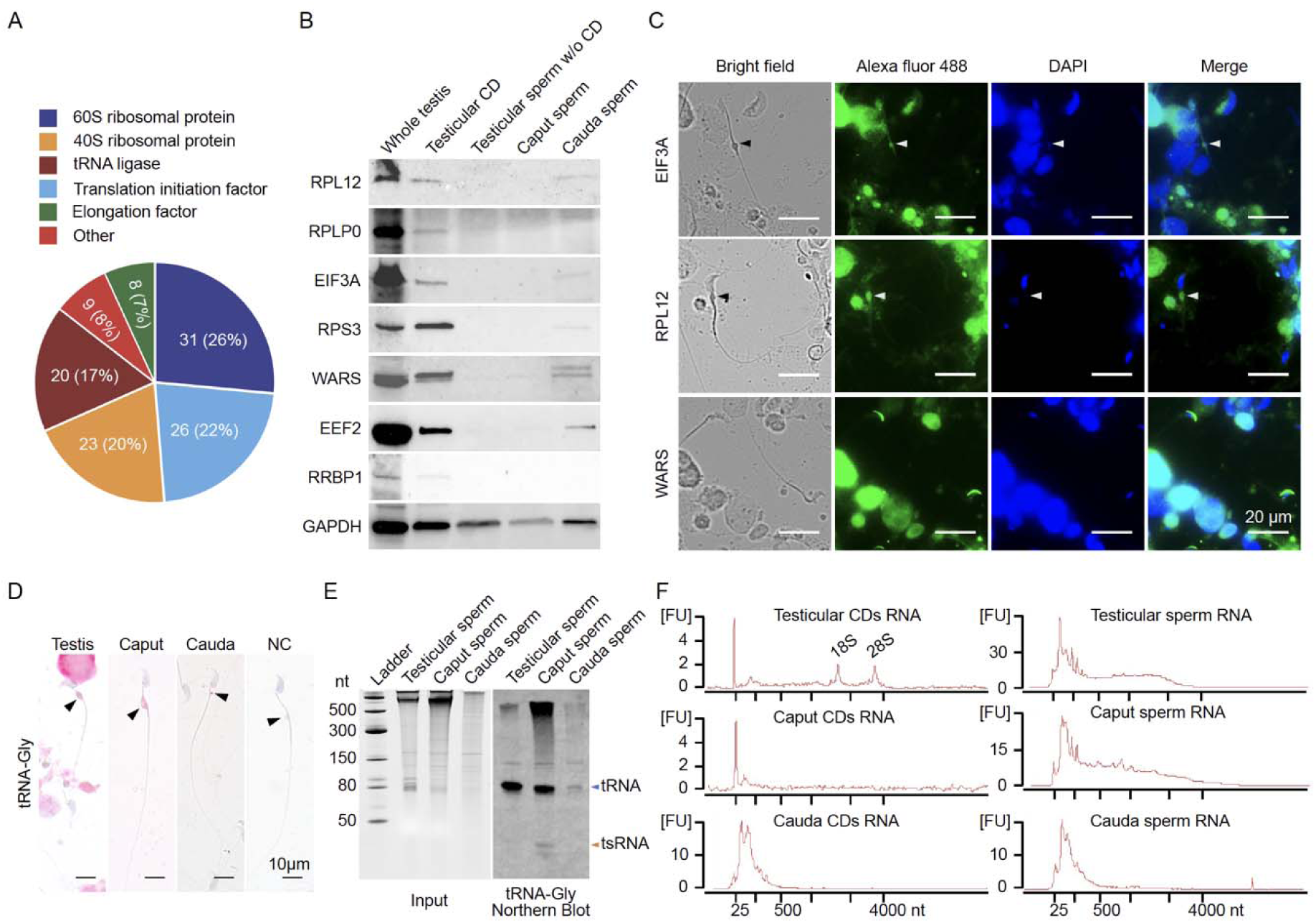
Testicular CDs contain intact cytoplasmic translational machinery. (A) Pie chart showing the composition of translational machinery proteins identified in CD proteomics, categorized by class. (B) Western blot analysis of translational machinery proteins in whole testis (positive control), testicular CDs, testicular sperm without CDs, caput sperm, and cauda sperm. GAPDH, loading control. (C) Immunofluorescence of EIF3A, RPL12, and WARS in testicular sperm. Scale bar = 20 μm. (D) Small RNA *in situ* hybridization (sRNA-ISH) of tRNA-Gly in testicular, caput, and cauda sperm. Arrowheads indicate CDs. Scale bar = 10 μm. (E) Northern blot of tRNA-Gly in testicular, caput, and cauda sperm, showing intact tRNA in testicular sperm and fragmentation in caput and cauda sperm. (F) Bioanalyzer RNA profiles of CDs and CD-less sperm from testis, caput, and cauda epididymis, demonstrating the presence of 28S and 18S rRNA exclusively in testicular CDs.

We next examined whether intact tRNAs and rRNAs, essential substrates for cytoplasmic translation, are preserved in testicular CDs. Small RNA *in situ* hybridization (ISH) showed that tRNA-Gly localizes predominantly to CDs of testicular, caput, and cauda sperm (Figure 3D). Northern blot analysis revealed that tRNA-Gly remains intact in testicular sperm but becomes fragmented upon entry into the caput epididymis (Figure 3E), coinciding precisely with the loss of cytoplasmic translational activity. This fragmentation is consistent with our previous observation that tRNA-derived small RNAs (tsRNAs) are generated from tRNAs during sperm transit in the caput epididymis^10^.

Bioanalyzer analysis of RNA from purified CDs and CD-less sperm demonstrated that 28S and 18S rRNAs are clearly present in testicular CDs, but absent from caput and cauda CDs and from CD-less sperm (Figures 3F and S4A). Collectively, these data demonstrate that intact cytoplasmic translational machinery indeed exists exclusively in testicular CDs.

### Ribo-lite identifies a translatome in testicular CDs enriched for spermatid development, cytoplasmic translation, energy production, and RNA metabolism

To capture the mRNAs actively engaged by ribosomes in testicular CDs, we applied Ribo-lite^39^, which profiles ribosome-protected fragments (RPFs) from cytoplasmic ribosomes using limited starting material (Figure 4A). Testicular, caput, and cauda CDs were treated with CHX to freeze ribosomes on their mRNAs and processed for Ribo-lite. RPFs were detected exclusively in testicular CDs, not in caput or cauda CDs (Figures 4B and S4B), confirming that active cytoplasmic ribosome engagement is restricted to the testicular stage. Testicular pellet (enriched for spermatocytes and round spermatids) and caput pellet (enriched for epididymal epithelial cells) were included as controls for contamination assessment (Figure 4B, S4B, S4C; Table S4).

**Figure 4.**
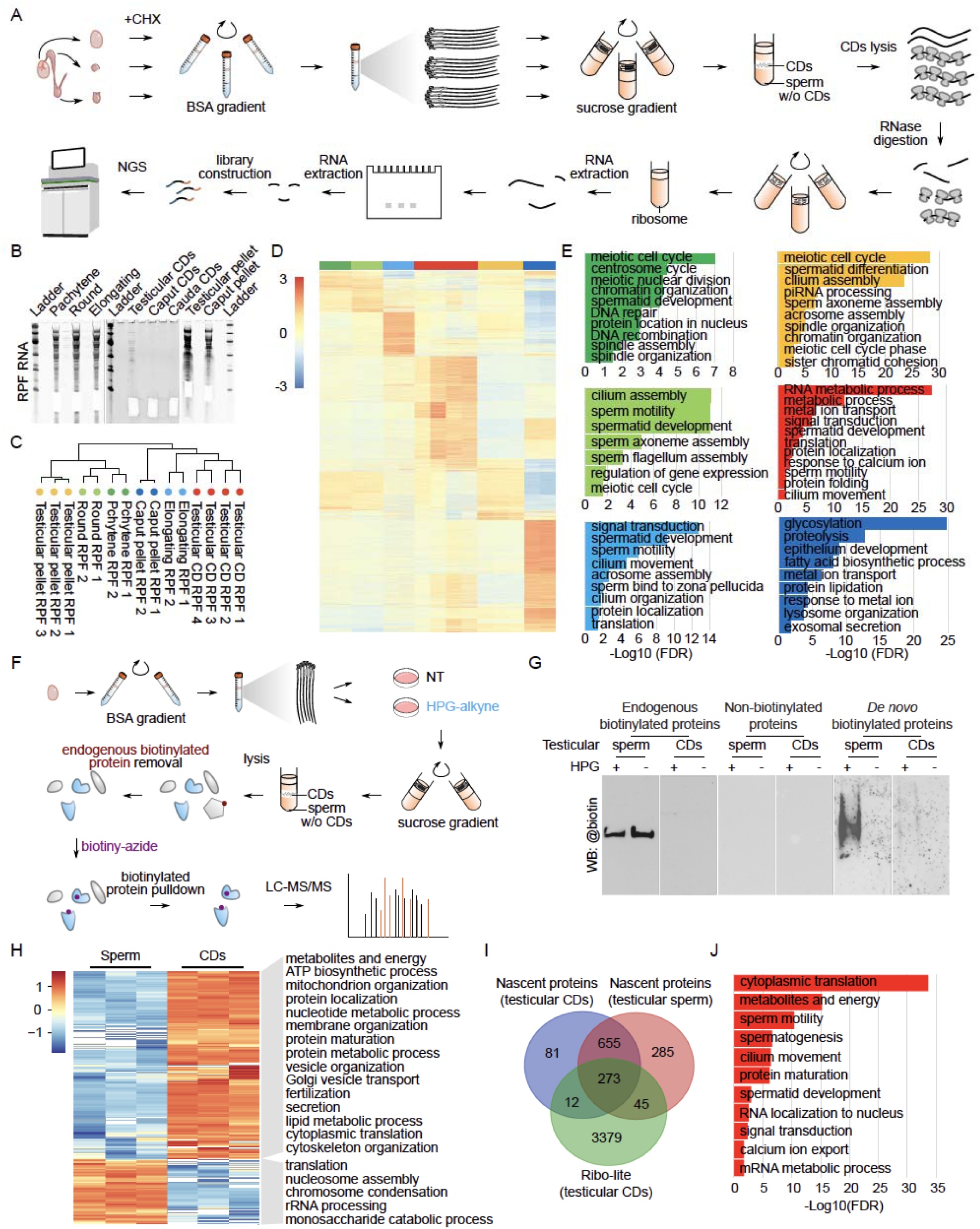
Ribo-lite and BONCAT identify the translatome and nascent proteome of testicular CDs, revealing enrichment for cytoplasmic translation, spermatid development, energy production, and RNA metabolism. (A) Workflow of Ribo-lite for testicular, caput, and cauda CDs. (B) Representative gel image of ribosome-protected fragments (RPFs) from pachytene spermatocytes, round spermatids, elongating spermatids, testicular CDs, caput CDs, cauda CDs, testicular pellets, and caput pellets. (C) Hierarchical clustering of translatome relationships across all cell types and CD populations. (D) Heatmap of normalized Ribo-lite data across all samples. Color codes as in (C). (E) GO term enrichments for each sample group. Color codes as in (C). (F) Workflow for enrichment of *de novo* synthesized proteins in testicular CDs and sperm using BONCAT. (G) Western blot showing biotinylated proteins (endogenous and nascent) at each stage of the BONCAT procedure. (H) Heatmap and GO term enrichment of differentially enriched nascent proteins in testicular CDs and sperm. (I) Venn diagram showing overlap between the Ribo-lite- identified translatome in testicular CDs and the BONCAT-identified nascent proteome in testicular sperm and CDs. (J) GO term enrichment of the intersection set shared among Ribo-lite RPFs and BONCAT nascent proteins.

Hierarchical clustering revealed that testicular CDs are clearly separated from testicular pellets (Figure 4C; Table S4), ruling out cross-contamination. To determine the cellular origin of CD-associated mRNAs, we performed Ribo-lite on purified pachytene spermatocytes, round spermatids, and elongating spermatids. Testicular CD RPFs clustered most closely with elongating spermatids (Figures 4C and 4D), indicating that the mRNAs being translated in testicular CDs originated from their immediate precursor cells. This is consistent with the developmental continuity between elongating spermatids and testicular sperm.

GO term enrichment analysis of the testicular CD translatome revealed overrepresentation of terms including spermatogenesis, spermatid development, sperm motility, RNA metabolic process, and cytoplasmic translation (Figure 4E; Table S5). By contrast, the testicular pellet translatome reflected the spermatocyte and round spermatid populations it contains, with enrichment for terms related to the meiotic cell cycle, DNA repair, P granule organization, and piRNA processing. The elongating spermatid translatome was enriched for acrosome assembly, sperm motility, spermatid development, cilium organization, and translation. Together, these results indicate that CDs actively translate a specific set of mRNAs inherited from elongating spermatids and centered on functions critical for sperm maturation.

### BONCAT identifies nascent proteins in testicular CDs involved in cytoplasmic translation, spermatid development, energy production and RNA metabolism

To directly capture newly synthesized proteins in testicular CDs and sperm, we adapted the BioOrthogonal Non-Canonical Amino Acid Tagging (BONCAT) approach^40,41^ and coupled it with LC-MS/MS proteomics (Figure 4F). Because sperm contain abundant endogenous biotinylated proteins that could confound streptavidin-based enrichment (Figure 4G), we developed a two-step protocol: first depleting endogenous biotinylated proteins using streptavidin beads, then labeling HPG-tagged nascent proteins with biotin-azide *via* click chemistry and re-enriching on streptavidin beads (Figure 4F). The efficiency of this depletion step was validated by Western blot (Figures 4G and S5A). HPG incorporation was confirmed to occur only in HPG-treated samples (Figure 4G).

BONCAT-LC-MS/MS identified 1,314 and 1,075 nascent proteins in testicular sperm and CDs, respectively (Table S6). After excluding proteins present in no-HPG controls, 1,263 sperm and 1,026 CD nascent proteins were retained (Figures 4H, S5B and S5C; Table S6). PCA confirmed that testicular sperm and CDs are distinct nascent proteomic populations (Figure S5D). Of these, 928 proteins were shared between sperm (∼73%) and CDs (∼90%) (Figure 4I), suggesting that the majority of newly synthesized proteins are shuttled from CDs into sperm after synthesis. GO analysis of CD-specific nascent proteins revealed enrichment for cytoplasmic translation, energy production, RNA metabolic process, and vesicle organization, whereas sperm-specific nascent proteins were enriched for chromosome condensation and RNA processing (Figures 4H and S5E; Table S7). ∼27% of CD nascent proteins and 22% of sperm nascent proteins were also present in the Ribo-lite-identified RPFs (Figure 4I), and this overlapping set was enriched for cytoplasmic translation, spermatogenesis, spermatid development, energy production, and mRNA metabolic process (Figure 4J; Table S7). These results confirm that testicular CDs are active sites of nascent protein synthesis and that the majority of newly made proteins are shared with sperm, likely reflecting transfer of CD-synthesized proteins into the sperm body.

### Cauda sperm possess intact mitochondrial translational machinery

Having established cytoplasmic translation in testicular CDs and its cessation in caput sperm, we turned to the mechanism of translational resumption in cauda sperm. Western blot demonstrated expression of mitochondrial ribosomal proteins MRPS16 and MRPL37 in cauda sperm (Figure 5A). Immunofluorescent staining revealed distinct localization patterns: MRPS16 was enriched at the sperm acrosome and midpiece across all three sperm populations, while MRPL37 was highly concentrated in CDs of testicular sperm and then redistributed to the midpiece in caput and cauda sperm (Figure 5B), suggesting that mitochondrial ribosomal components undergo dynamic spatial reorganization during epididymal maturation. Northern blot detected strong signals for the mitochondrial rRNAs *mt-Rnr1* and *mt-Rnr2* in cauda sperm (Figure 5C).

**Figure 5.**
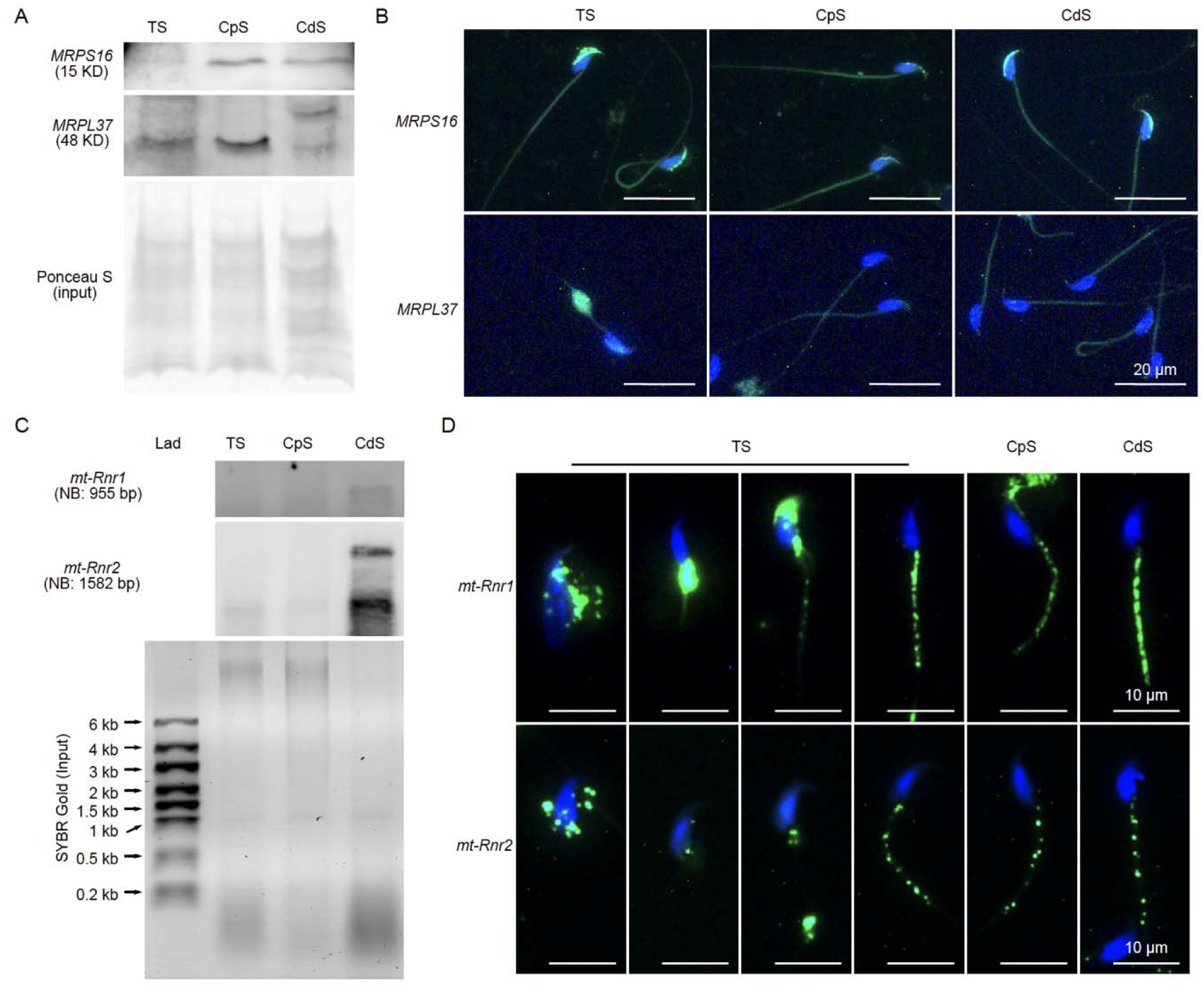
Cauda sperm possess intact mitochondrial translational machinery. (A) Western blot of mitochondrial ribosomal proteins MRPS16 and MRPL37 in testicular sperm (TS), caput sperm (CpS), and cauda sperm (CdS). (B) Immunofluorescence of MRPS16 and MRPL37 in TS, CpS, and CdS. Scale bar = 20 μm. (C) Northern blot of mitochondrial rRNAs *mt-Rnr1* and *mt-Rnr2* in TS, CpS, and CdS. (D) RNA FISH of *mt- Rnr1* and *mt-Rnr2* in TS, CpS, and CdS. Scale bar = 10 μm. Note punctate midpiece signals in cauda sperm, indicating translational activity in a subpopulation of midpiece mitochondria.

RNA FISH showed that both transcripts localize to the midpiece of caput and cauda sperm, with fewer signals in caput sperm consistent with ongoing midpiece maturation^42^. In testicular sperm, *mt-Rnr1* and *mt-Rnr2* showed a more dispersed distribution, progressively concentrating at the midpiece as sperm mature (Figure 5D). Notably, both mt-rRNAs displayed punctate, dashed-line signals along the sperm midpiece, indicating that only a subpopulation of mitochondria along the midpiece is translationally active (Figure 5D). Together, these data indicate that cauda sperm contain the essential protein and RNA components of the mitochondrial translational machinery.

### Cauda sperm synthesize nascent proteins involved in mitochondrial function, calcium signaling, and flagellum assembly

To identify the proteins being *de novo* synthesized by cauda sperm mitochondrial ribosomes, we applied a modified SUnSET strategy using 5’-azido-puromycin (AP) followed by DBCO-PEG4-Biotin click chemistry and streptavidin enrichment for LC-MS/MS (Figures 6A, S5F and S5G, Table S8). PCA confirmed that AP-treated (PB), untreated (NT), and their respective input samples are distinct populations (Figure 6B). The PB condition enriched 84 proteins relative to PB-input (Figure 6C), of which 38 remained after excluding those also present in NT or NT-input controls (Figures 6D and S5H). These 38 *de novo* synthesized proteins fall into nine functional categories (Figures 6E and 6F): mitochondrial function (39%; including mt-ND5, COX20, SERAC1^43,44^, NIT1^45^, and ATP6V1H^46^, calcium channels (13%; including CATSPERD^47^, S100A14^48^, and STIM1^49^, and flagellum assembly (13%; including MYCBPAP ^50^, CCDC159 ^51^, and PPIL6^52^, with additional proteins in acrosomal maturation, chromatin condensation, spermatogenesis, epididymis-related functions, and cytoplasmic translation.

**Figure 6.**
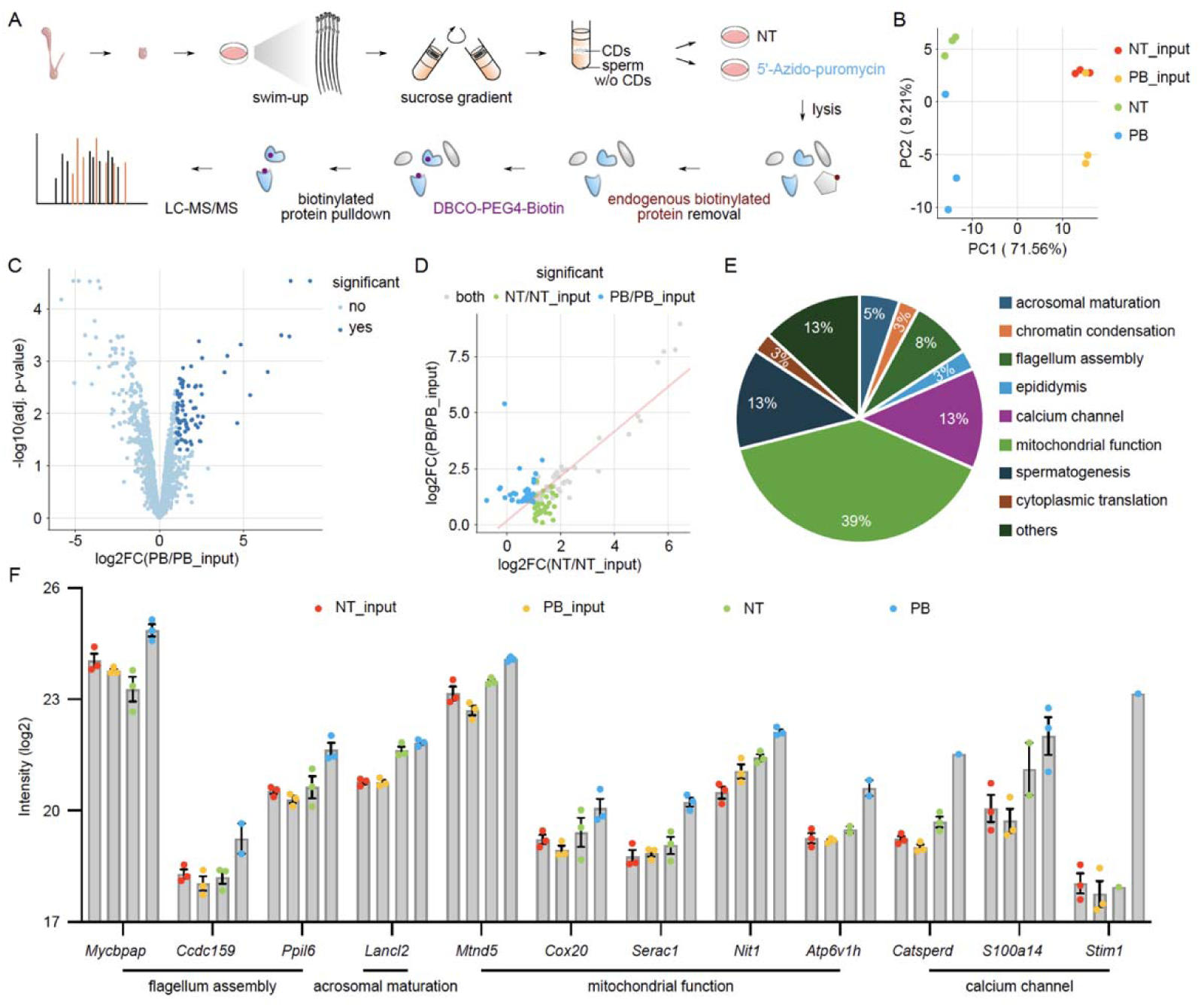
Cauda sperm synthesize nascent proteins involved in mitochondrial function, calcium signaling, and flagellum assembly. (A) Workflow for enrichment of *de novo* synthesized proteins in cauda sperm using the SUnSET approach with 5’-azido-puromycin (AP). (B) PCA of cauda sperm puromycin- biotin (PB), no-treatment (NT), PB-input, and NT-input proteomics data. (C) Volcano plot of PB-enriched proteins relative to PB-input. (D) Fold-change plot comparing PB/PB-input versus NT/NT-input enrichment, showing PB-specific proteins. (E) Pie chart of functional categories of PB-specific *de novo* synthesized proteins. (F) Bar graph showing protein intensities across PB, NT, PB-input, and NT-input for selected PB-specific proteins, grouped by functional category.

Western blot confirmed upregulation of S100A14, ATXN3^53^, and mt-ND5 in cauda sperm (Figure S5I). Immunofluorescence revealed dynamic spatial redistribution of these proteins: in testicular sperm, they surround the sperm head and reside within CDs, then progressively concentrate at the sperm midpiece in caput and cauda sperm with increasing intensity (Figure S5J). NIT1 followed a similar but distinct trajectory, accumulating at the connecting piece in caput and cauda sperm (Figure S5J). SERAC1 localized to CDs in testicular sperm, shifted to the acrosome in caput sperm, and then concentrated in both the sperm head and midpiece in cauda sperm (Figure S5J). These findings indicate that cauda sperm synthesize a specific set of proteins *via* mitochondrial translation that equip them to meet the functional demands of fertilization, including ATP production, calcium-triggered hyperactivation, and flagellar mechanics.

### Cytoplasmic and mitochondrial translation are each causally required for sperm maturation and function

To establish a direct causal link between CD-based cytoplasmic translation and sperm functional competence *in vivo*, we performed intra-rete testis injection of cycloheximide (CHX) or DMSO vehicle into adult male mice (Figure 7A). Injection volume was restricted to <2 μL to confine inhibition to the rete testis (a reservoir of testicular sperm) without backflowing into the seminiferous tubules, which would potentially disrupt spermatogenesis. Consistent with this specificity, testicular histology was indistinguishable between CHX- and DMSO-injected groups (Figure 7B), and sperm morphology was similarly unaffected (Figures 7C and 7D). However, four days post-injection, the time required for sperm transit from testis to cauda^6^, CHX-treated mice exhibited a significantly reduced concentration of swim-out sperm and a pronounced decrease in progressive motility compared to DMSO controls (Figures 7E and 7F), demonstrating that cytoplasmic translation in testicular sperm is causally required for the acquisition of normal sperm motility.

**Figure 7.**
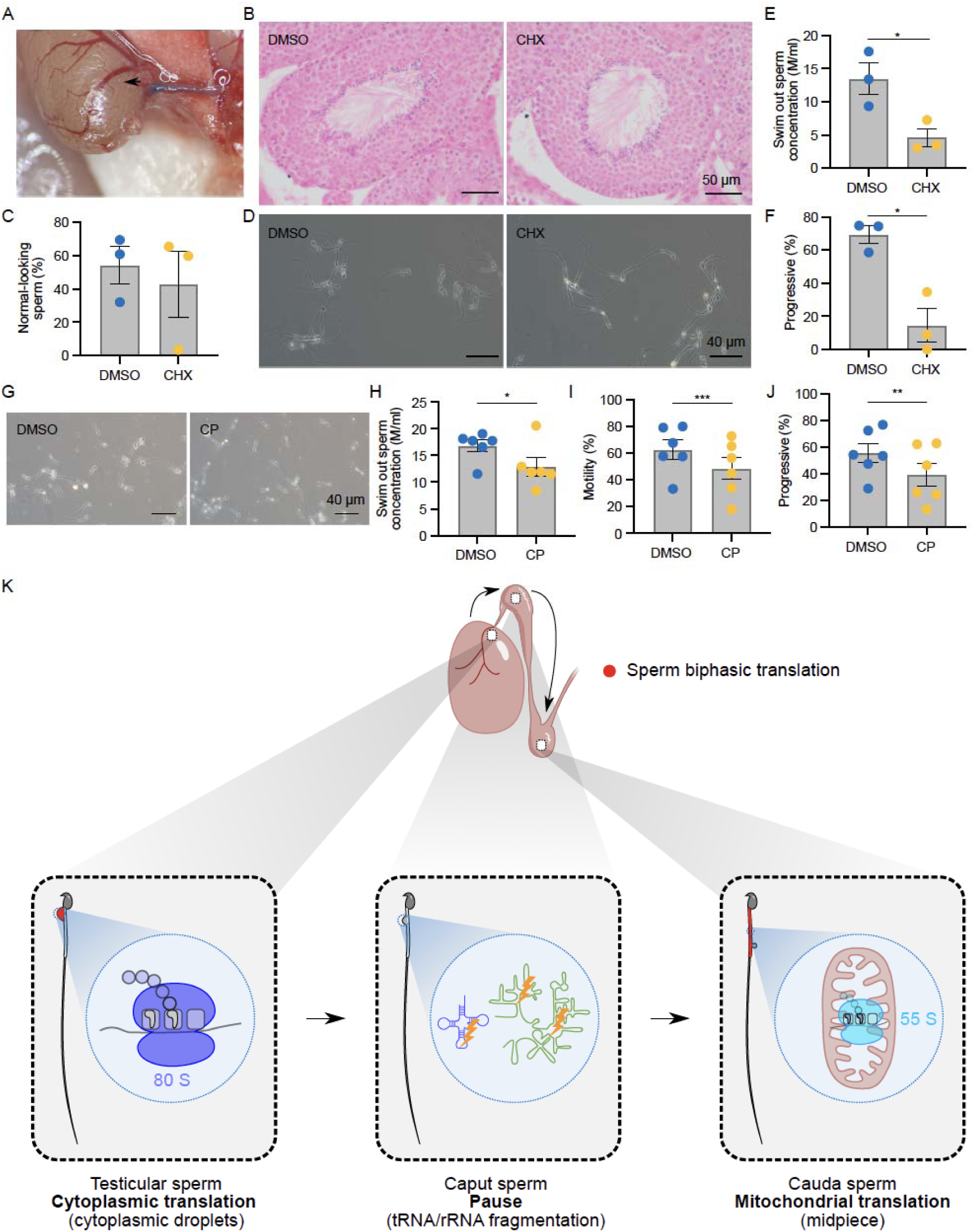
Cytoplasmic and mitochondrial translation are required for sperm maturation and function. (A) Representative image of intra-rete testis injection. The arrowhead indicates the site of needle insertion. Trypan blue was used to confirm successful injection; only a light blue color was observed in a small subset of seminiferous tubules to minimize side effects on spermatogenesis. (B) Hematoxylin and eosin (H&E) staining of testis sections from DMSO- and CHX-injected mice. Scale bar = 50 μm. (C) Percentage of morphologically normal sperm after rete testis injection with DMSO or CHX (n.s., not significant). (D) Representative micrographs of sperm morphology 4 days after rete testis injection with DMSO or CHX. Scale bar = 40 μm. (E) Sperm concentration 4 days after rete testis injection with DMSO or CHX. *p < 0.05. (F) Progressive sperm motility 4 days after rete testis injection with DMSO or CHX. *p < 0.05. (G) Representative micrographs of cauda sperm morphology after treatment with DMSO or chloramphenicol (CP) during capacitation in HTF medium. Scale bar = 40 μm. (H) Sperm concentration after DMSO or CP treatment during capacitation. *p < 0.05. (I) Total sperm motility after DMSO or CP treatment. ***p < 0.001. (J) Progressive sperm motility after DMSO or CP treatment. **p < 0.01. Data are presented as mean ± SEM. (K) Schematic model of the biphasic translational program during sperm epididymal maturation. Testicular sperm retain a cytoplasmic droplet (CD) harboring complete cytoplasmic translational machinery that actively synthesizes nascent proteins (80S ribosomes, red dot). As sperm transit through the caput epididymis, tRNA fragmentation and tsRNA generation silence CD-based cytoplasmic translation. In cauda sperm, protein synthesis resumes via mitochondrial ribosomes (55S) localized to the midpiece, enabling transcriptionally silent sperm to maintain continuous translational capacity throughout epididymal maturation.

To test whether mitochondrial translation in cauda sperm is likewise required for function, cauda sperm were first treated with chloramphenicol (CP, a mitochondrial translation inhibitor) or DMSO vehicle, followed by HTF incubation for capacitation (Figure 7G). Sperm morphology was comparable between groups (Figure 7G), but CP-treated sperm showed significant reductions in swim-out sperm concentration, total motility, and progressive motility relative to DMSO controls (Figures 7H–7J). These results demonstrate that mitochondrial translation is essential for normal sperm capacitation and motility. Together, these pharmacological experiments establish that both phases of the biphasic translational program, cytoplasmic translation in testicular CDs and mitochondrial translation in cauda sperm, are causally required for the acquisition of motility and fertilization competence.

## Discussion

Spermatozoa have long been thought to be translationally inert on the grounds that transcriptional silencing during spermiogenesis eliminates the RNA-production capacity needed to replenish ribosomes and mRNAs. This assumption has been difficult to reconcile with the well-documented acquisition of new functional properties, motility, membrane remodeling, and zona-binding competence during the ∼10 days sperm spend traversing the epididymis. Our data resolve this paradox by uncovering a biphasic translational program that sustains protein synthesis throughout epididymal maturation despite transcriptional silence. Testicular sperm exploit a complete cytoplasmic translational machinery preserved within cytoplasmic droplets; this machinery is then switched off as sperm enter the caput epididymis, and translation resumes *via* mitochondrial ribosomes in cauda sperm. This sequential use of two mechanistically distinct translational systems represents an unanticipated solution to the challenge of protein homeostasis in a cell that can neither transcribe new RNAs nor simply maintain static protein inventories. A schematic summary of this biphasic translational program is presented in Figure 7K, which illustrates the three sequential translational states of maturing sperm: active cytoplasmic translation via 80S ribosomes housed in the CD of testicular sperm; translational silence in caput sperm coincident with tRNA fragmentation; and resumption of protein synthesis via mitochondrial 55S ribosomes localized to the midpiece of cauda sperm.

The identification of cytoplasmic droplets as functional translational organelles is, in retrospect, consistent with a body of structural evidence that has not previously been given a clear mechanistic interpretation. Ultrastructural studies conducted decades ago identified monosomes, polysomes, and Golgi-like membrane systems within CDs^19–21^. CD proteomics revealed a striking complement of ribosomal proteins and translation factors^12,14^. Our prior work showed that CDs contain not only mRNAs and circular RNAs with coding potential^22^, but also dynamic populations of tsRNAs and rsRNAs^10^, small RNAs now recognized to participate in translational regulation^27–32^. The present study ties these observations together by directly demonstrating that CDs harbor intact 18S and 28S rRNAs and tRNAs, express all major classes of cytoplasmic translation factors, and actively incorporate puromycin into nascent proteins in a CHX-sensitive, CP- insensitive manner. The fact that this activity is lost when CDs are removed from sperm and is absent from isolated caput and cauda CDs confirms that translational competence is a property of testicular CDs.

The Ribo-lite and BONCAT data together provide a detailed portrait of what testicular CDs translate and synthesize. The translatome, mRNAs protected by actively elongating ribosomes, is enriched for spermatid development, sperm motility, cytoplasmic translation, and RNA metabolic processes, reflecting a biosynthetic program focused on equipping sperm for their journey ahead. The clustering of testicular CD RPFs with elongating spermatid RPFs argues that CDs inherit their translational substrate pool from their immediate precursor cells; the mRNAs present in CDs are likely those being actively translated in elongating spermatids at the time of spermiation, and these are retained and continue to be translated in the CD after spermiation. This inheritance of translational state from elongating spermatids to testicular CDs represents a form of cytoplasmic memory, connecting the spermatids’ biosynthetic program to the early stages of sperm maturation. The fact that roughly 90% of CD-synthesized nascent proteins are also detected in testicular sperm suggests active protein transfer from CDs into the sperm body, a directional flux consistent with the known role of CDs as protein exporters during epididymal transit^9,12,14,17^.

A key question raised by this model is how cytoplasmic translation is terminated as sperm enter the caput epididymis. The coincidence of translational silencing with the fragmentation of tRNA into tsRNAs is unlikely to be accidental. tsRNAs derived from the 5’ end of tRNAs have been shown to suppress translation initiation by displacing eIF4G and eIF4A from mRNA caps and inhibiting ribosome assembly^28,30^. More broadly, 5’ tsRNAs can displace PABP from poly(A) tails and sequester translation machinery into stress granule-like structures^32^. In the context of sperm, the caput-specific generation of tsRNAs from full-length tRNAs^10^ would simultaneously eliminate the amino acid acceptor substrates needed for elongation and produce inhibitory small RNAs capable of blocking initiation. This dual mechanism, substrate depletion plus active suppression, may explain why translational shutdown is so complete in caput sperm. The upstream trigger for tRNA fragmentation itself remains to be identified, but epididymal luminal factors, epithelial-derived exosomes, or RNase activation represent plausible candidates. The loss of *Dnmt2*, an enzyme that methylates tRNA to protect it from cleavage, alters tsRNA profiles without affecting male fertility^54–56^, suggesting that the precise tsRNA species generated, rather than tRNA fragmentation *per se*, may be the relevant regulatory variable, and that cytoplasmic translation is dispensable by the time sperm enter the caput.

The resumption of translation in cauda sperm *via* mitochondrial ribosomes is equally striking, and our data reveal several features of this process that were not previously appreciated. The mitochondrial ribosomal proteins MRPS16 and MRPL37 are present in testicular sperm but reorganize during epididymal transit, with MRPL37 shifting from a CD localization in testicular sperm to the midpiece in caput and cauda sperm. The mitochondrial rRNAs *mt-Rnr1* and *mt-Rnr2* are similarly repositioned to the midpiece, but their punctate distribution along the midpiece suggests that not all mitochondria in cauda sperm are translationally active. This could reflect functional heterogeneity among the ∼70–75 midpiece mitochondria or indicate that mitochondrial translation is spatially restricted to specific mitochondrial compartments. The determinants of this heterogeneity remain to be established, but candidate mechanisms include variation in membrane potential along the midpiece, unequal mtDNA copy number among individual mitochondria, or positional cues imposed by the fibrous sheath and outer dense fibers that could restrict ribosome assembly or access to mitochondrial transcripts at specific axonemal segments. A prior study demonstrated that sperm mitochondria are immature at the time of testicular release and require epididymal transit for functional maturation^42^, which is consistent with our observation that mitochondrial translational machinery becomes midpiece-organized only in cauda sperm. The nature of the signal that triggers this maturation and coordinates the handoff from cytoplasmic to mitochondrial translation remains an important open question.

The identities of the proteins synthesized by cauda sperm mitochondrial ribosomes provide insight into the functional priorities of this translational phase. The largest category (39%) comprises mitochondrial function proteins, including mt-ND5 (a complex I subunit), COX20 (a complex IV assembly factor), and SERAC1 (required for mitochondrial membrane remodeling and mtDNA maintenance)^43,44^. Our finding that sperm synthesize core respiratory chain components in the cauda epididymis suggests that oxidative phosphorylation capacity is still being established or maintained at this stage, consistent with the increasing energetic demands of motility acquisition. The next largest categories, calcium channels (13%; CATSPERD, S100A14, STIM1) and flagellum assembly proteins (13%; MYCBPAP, CCDC159, PPIL6), reflect two other critical dimensions of sperm fertilization competence: the calcium signaling required for hyperactivation and chemotaxis toward the egg^47–49^, and the flagellar mechanics required for penetration of the zona pellucida^50–52^. The fact that all three functional categories are represented in the nascent proteome of CD-less cauda epididymal sperm suggests that mitochondrial translation in cauda sperm is not merely maintaining cellular homeostasis but actively building the molecular toolkit required for fertilization.

The functional necessity of each translational phase is directly established by pharmacological intervention. CD-based cytoplasmic translation is causally required for sperm motility: intra-rete testis injection of cycloheximide, which blocks cytoplasmic ribosomes in translationally active CDs, almost completely abolishes sperm motility.

Mitochondrial translation in cauda sperm is equally indispensable: pre-treatment of mature cauda sperm with chloramphenicol before HTF-induced activation abolishes hyperactivated motility, the high-amplitude asymmetric flagellar beating that is mechanistically required for zona pellucida penetration and fertilization. Together, these two experiments establish that neither translational phase is dispensable and that each contributes distinct functional outputs: acquisition of basal motility from CD-based cytoplasmic translation and capacity for hyperactivation from mitochondrial translation, both of which are necessary for fertilization competence. This result argues that the proteins synthesized by CDs during and shortly after spermiation are not redundant with pre-existing sperm proteins but are functionally indispensable for a key sperm property.

The specificity of the motility defect is particularly telling: it implies that among the large ensemble of proteins synthesized by testicular CDs, a subset is required for the normal acquisition or maintenance of motility. Whether this reflects synthesis of motility apparatus components, energy metabolism enzymes, signaling molecules, or RNA-binding proteins that coordinate downstream events in the epididymis will be an important focus of future work.

Viewed in a broader context, the biphasic translational program we describe here adds an unexpected dimension to the biology of post-meiotic male germ cells. It has long been appreciated that haploid spermatids address the problem of phase-specific protein production through translational delay: they transcribe mRNAs during the diploid spermatocyte stage, store them in a repressed form, and translate them at appropriate times during spermiogenesis^3^. What was not appreciated is that this translational capacity extends beyond spermiogenesis to post-testicular maturation, with CDs serving as a vehicle to carry active translational machinery out of the testis. In effect, the CD represents a cytoplasmic organelle that the elongated spermatid loads with translational machinery and substrate mRNAs before spermiation, and that sperm then utilize during the early epididymal phase before switching to mitochondrial translation for the remainder of maturation. This organization may reflect an evolutionary solution to the challenge of completing sperm maturation in the epididymal lumen, an environment where sperm are physically isolated from Sertoli cell support and must rely on their own biosynthetic resources.

Several important questions remain to be addressed. The mechanism by which cytoplasmic translation in CDs is terminated remains incompletely understood, though tRNA fragmentation and tsRNA-mediated suppression are the most parsimonious candidates. The identity of the signals that promote mitochondrial translational maturation in cauda sperm, and whether these involve epididymal epithelial-derived factors delivered *via* exosomes or other secretory mechanisms, is unknown. Whether the biphasic translational program we describe in mice is conserved in human sperm, where CDs are also present and are likewise considered markers of defective maturation when retained in ejaculated sperm^13^, has direct clinical implications for understanding male infertility. The observation that persistent CDs in ejaculated sperm are associated with poor outcomes^13^ suggests that the normal program of CD-based translation and eventual droplet loss is diagnostically and functionally relevant to human fertility.

In summary, we have uncovered a previously unrecognized mechanism by which sperm maintain translational competence during epididymal maturation. By sequentially exploiting cytoplasmic translation in transient organelles and mitochondrial translation in a permanent organelle, sperm achieve continuous protein synthesis across a developmental journey defined by transcriptional silence. CDs are not passive cytoplasmic remnants awaiting disposal, but active biosynthetic organelles whose output is causally required for normal sperm function. These findings open new avenues for understanding the molecular basis of epididymal sperm maturation and for identifying translational targets relevant to male infertility.

### Limitations of the study

Several limitations of the present study merit consideration. First, although our data implicate tRNA fragmentation and tsRNA-mediated translational suppression as the mechanism underlying translational silencing in caput sperm, direct functional evidence that specific tsRNA species are sufficient to silence translation in this context is lacking; the upstream signals that trigger tRNA cleavage during the testis-to-caput transition remain unknown. Second, all experiments were performed in mice, and whether the biphasic translational program described here, cytoplasmic translation in CDs followed by mitochondrial translation in the cauda, is conserved in human sperm has not been directly demonstrated; given the anatomical and physiological differences between rodent and human epididymides, extrapolation should be made with caution. Third, our imaging data indicate that only a subpopulation of midpiece mitochondria in cauda sperm are translationally active, yet the determinants of this heterogeneity, whether related to membrane potential, local mtDNA copy number, or structural constraints imposed by the axoneme, remain unresolved. Finally, the full repertoire of proteins synthesized by CDs and by mitochondrial ribosomes in cauda sperm, and their relative contributions to sperm function in vivo, await comprehensive proteomic characterization.

## Acknowledgments

The authors thank Rebekah J. Woolsey and David R. Quilici at the Nevada Proteomics Center, and Guihua (Eileen) Yue and John Clarke at the WSU Proteomics Service Center for their assistance with proteomic analyses. This work was supported by grants from the NIH/Eunice Kennedy Shriver National Institute of Child Health and Human Development (NICHD) (HD071736, HD085506, HD098593, HD099924, and P30GM110767 to W.Y.), NIH/National Center for Advancing Translational Science (NCATS) UCLA CTSI (UL1TR001881-01 to Z.W. and W.Y.), John Templeton Foundation (PID: 61174 to W.Y.), California Institute for Regenerative Medicine (EDUC4-12837 to Z.W. and W.Y.), Washington State University startup (PG00023119 to W.Y.).

## Author Contributions

Conceptualization, W.Y.; Methodology, Z.W., H.W., S.M., S.Chen, S.Crane, D.M., H.M., A.E.Y., S.K., and B.N.; Software, Z.W., R.D.M.M.; Validation, Z.W., H.W., S.Chen, S.M., D.M., and R.D.M.M.; Formal Analysis, Z.W., H.W., R.D.M.M.; Investigation, Z.W., H.W., and W.Y.; Resources, Z.W., H.W., S.Chen, and H.Z.; Data Curation, Z.W., H.W., and R.D.M.M.; Writing – Original Draft, Z.W., H.W., and W.Y.; Writing – Review & Editing, Z.W., H.W., and W.Y.; Visualization, Z.W., H.W., and R.D.M.M.; Supervision, Z.W. and W.Y.; Project Administration, Z.W. and W.Y.; Funding Acquisition, Z.W., H.W., and W.Y.

## Declaration of Interests

The authors declare no competing interests.

## STAR METHODS

## KEY RESOURCES TABLE

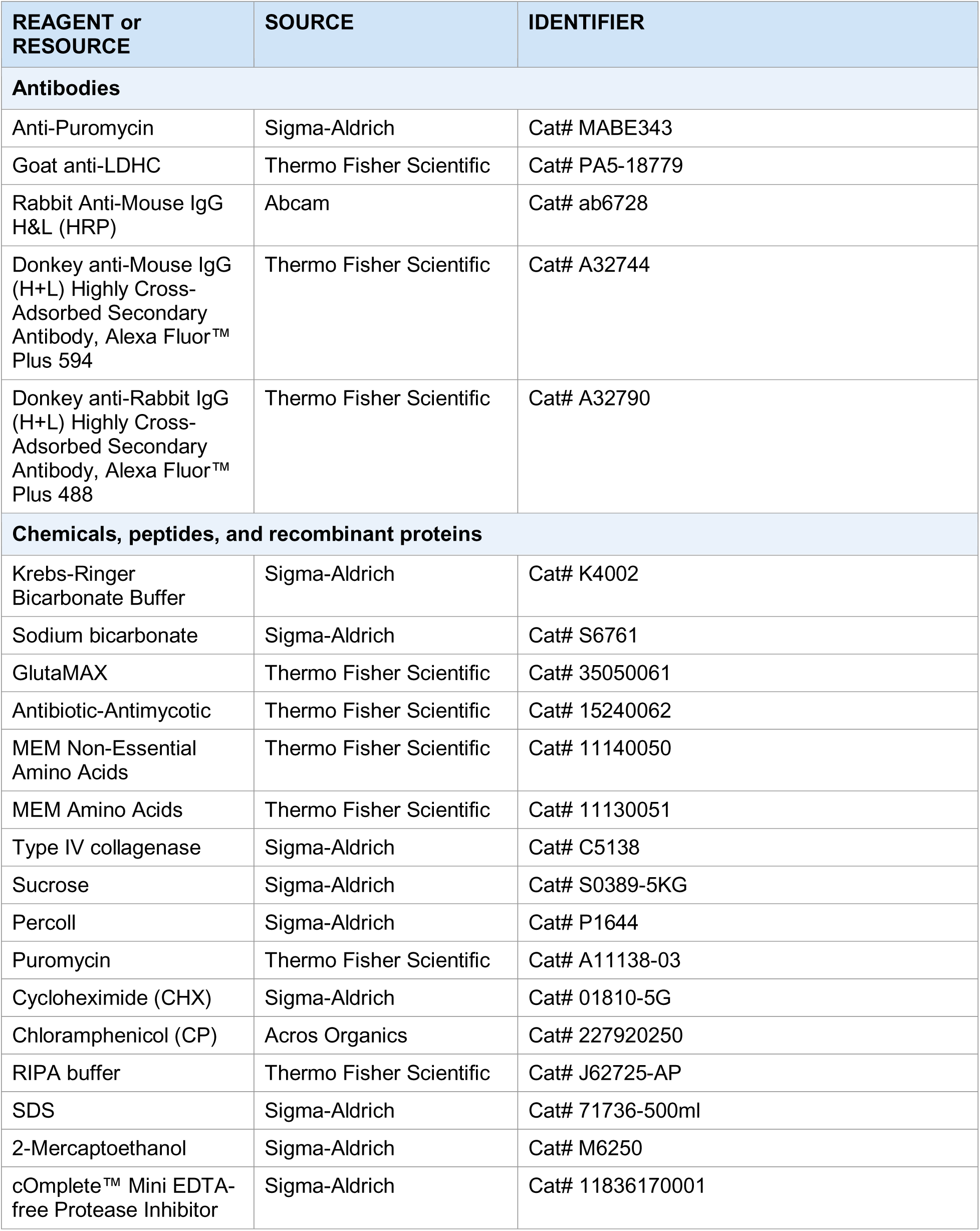
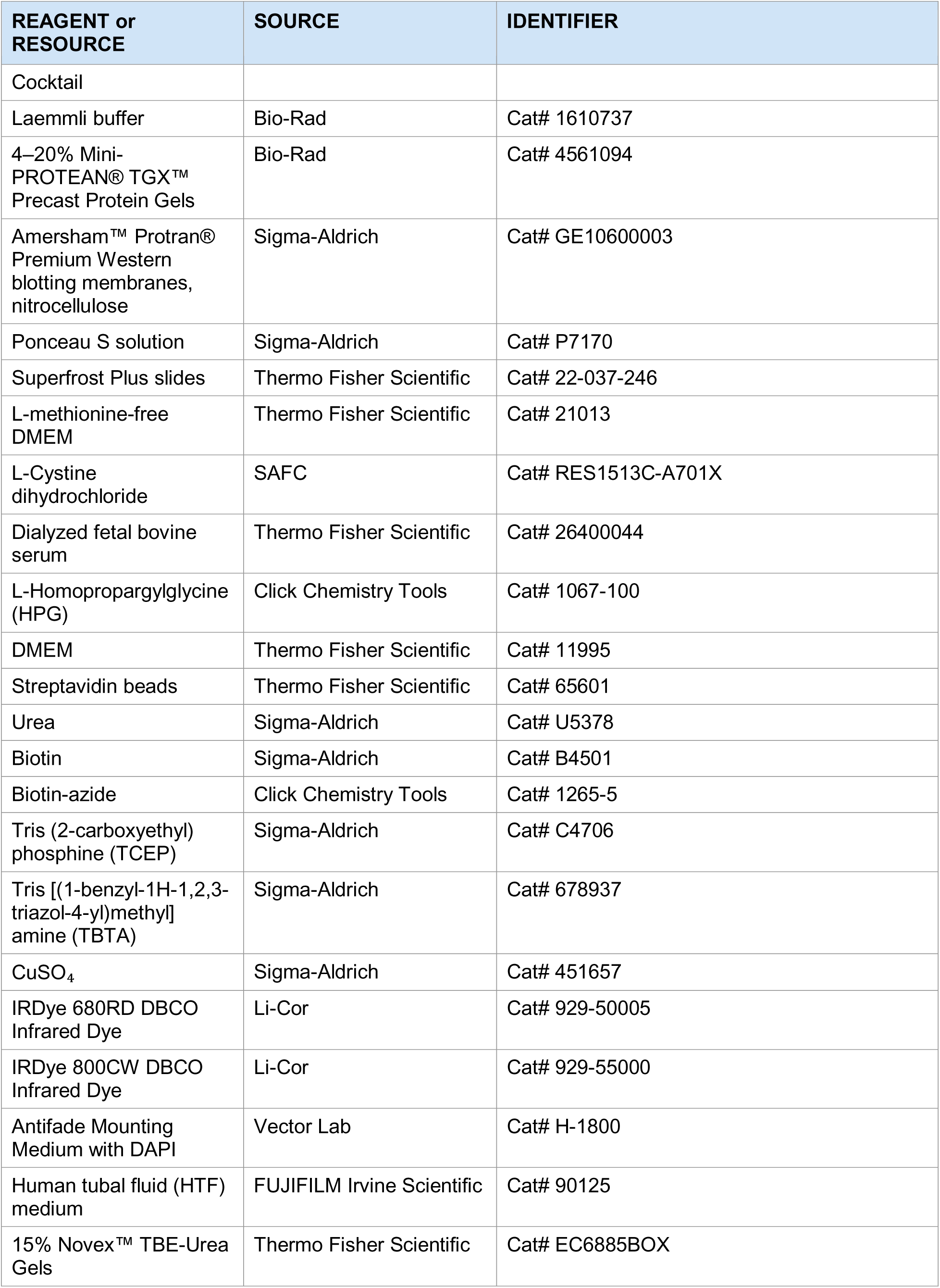
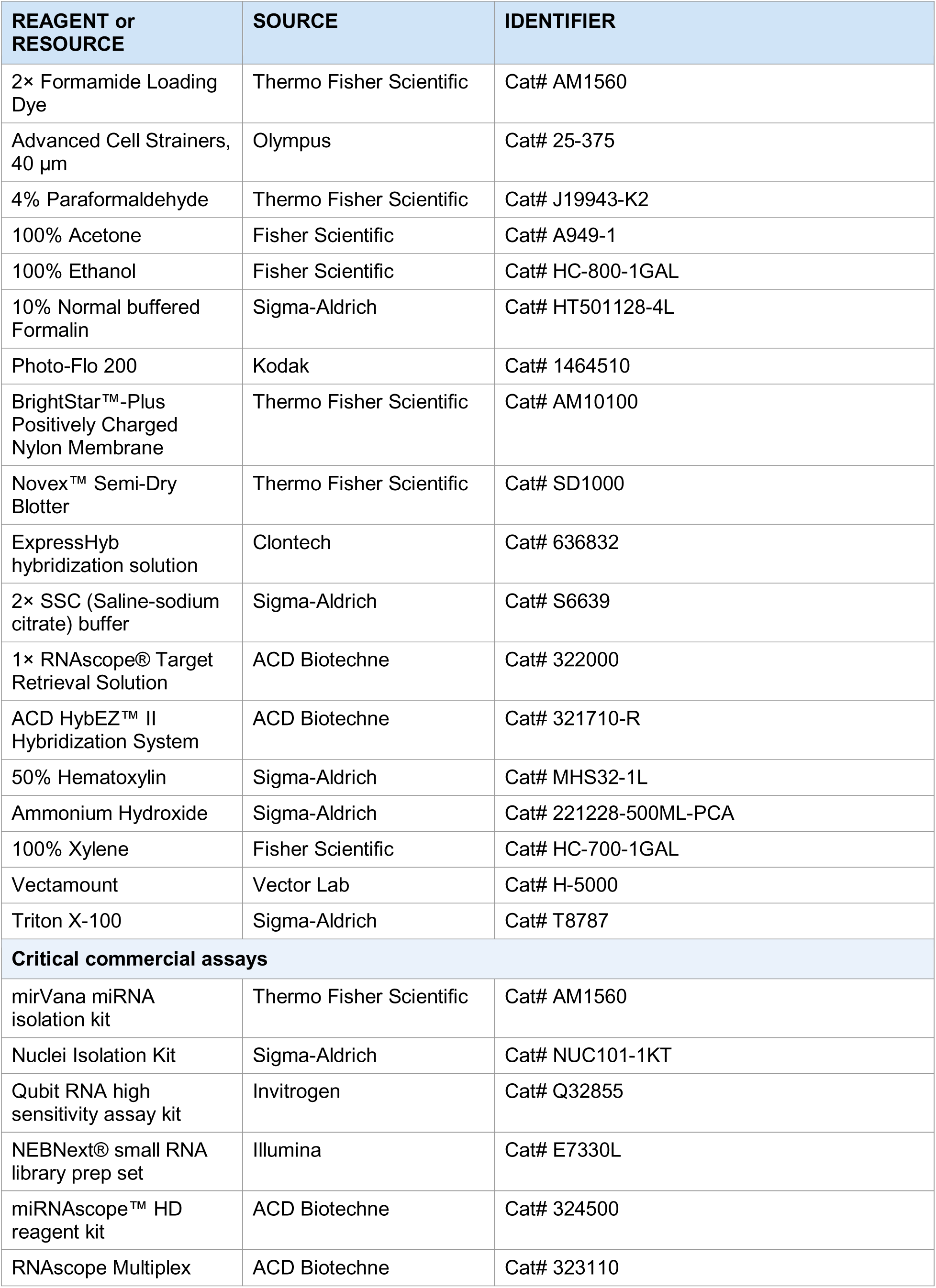
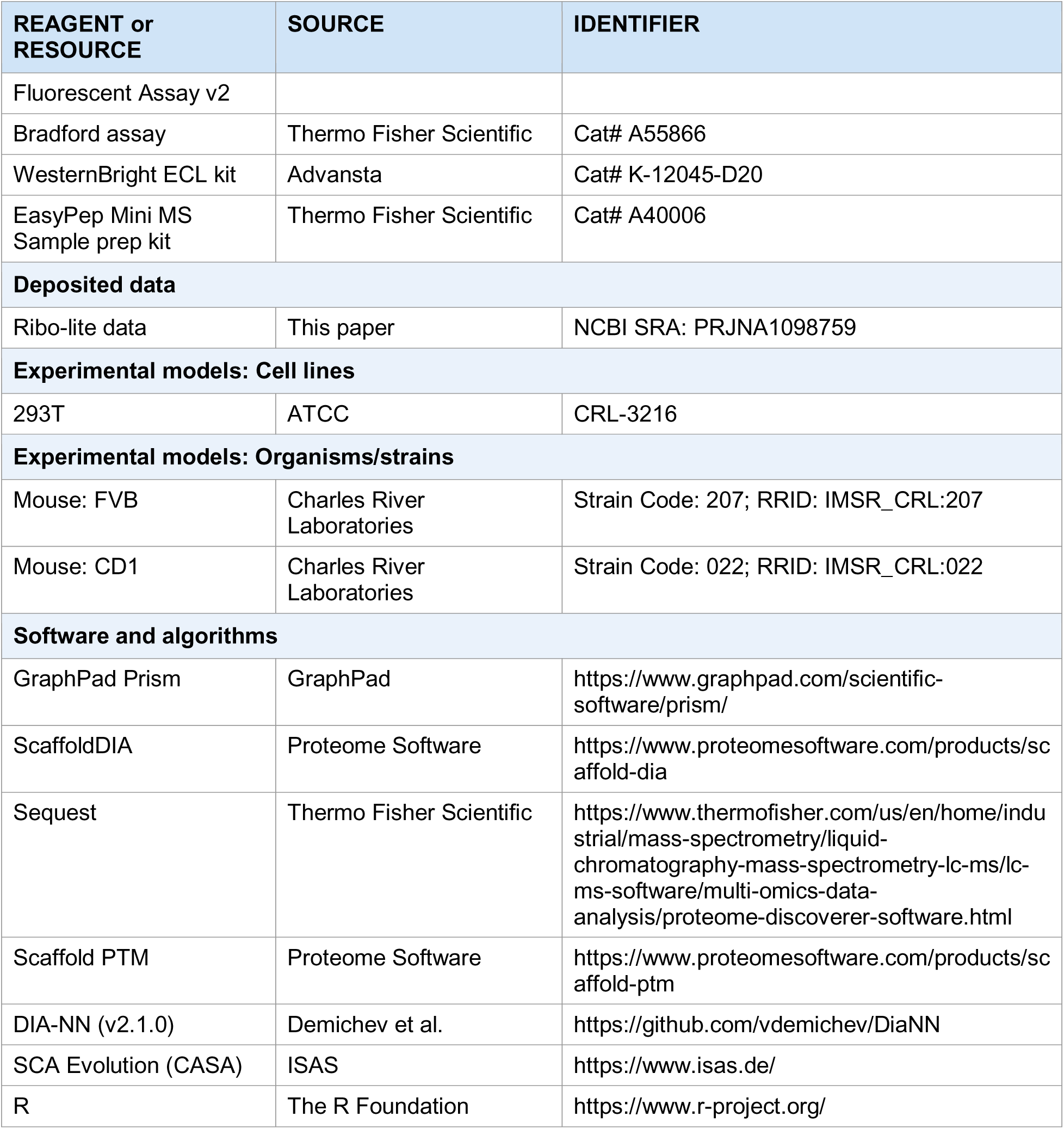

### RESOURCE AVAILABILITY

#### Lead contact

Further information and requests for resources and reagents should be directed to and will be fulfilled by the lead contact, Wei Yan.

#### Materials availability

Unique materials generated in this study are available from the lead contact without restriction.

#### Data and code availability

The Ribo-lite data have been deposited into the National Center for Biotechnology Information Sequence Read Archive database (accession no. PRJNA1098759). The mass spectrometry proteomics data have been deposited to the ProteomeXchange Consortium *via* the PRIDE partner repository with the dataset identifier PXD079797 and project DOI:10.6019/PXD079797 (Reviewer access details: Log in to the PRIDE website using the following details: Project accession: PXD079797; Token: tJ3voe3UmD36). Any additional information relating to the data reported in this paper is available from the lead contact upon request.

### EXPERIMENTAL MODEL AND SUBJECT DETAILS

#### Laboratory mice

All mice used in this study were ∼3-month-old adults on either a CD1 background (Strain #022, Charles River Laboratories, RRID:IMSR_CRL:022) or FVB background (Strain #207, Charles River Laboratories, RRID:IMSR_CRL:207). Animals were housed in a temperature- and humidity-controlled, specific pathogen-free facility under a 12:12- hour light-dark cycle with food and water ad libitum. All animal use protocols were approved by the Institutional Animal Care and Use Committees (IACUC) of the Lundquist Institute at Harbor-UCLA (protocol: 32132-03) and Washington State University (ASAF# 7444 and 7433), and all procedures followed the “Guide for the Care and Use of Laboratory Animals” established by the National Institutes of Health (1996, revised 2011).

### METHOD DETAILS

#### Pachytene spermatocytes, round and elongating spermatids, testicular, caput, and cauda sperm purification

Pachytene spermatocytes, round and elongating spermatids were purified as previously described^57–59^. At least four mice were used per sperm collection. Mice were anesthetized by intraperitoneal injection of avertin working solution (20 mg/mL in 0.9% saline; 0.8 mL per 20 g body weight) through a 25G needle (Cat# 309626, BD). Once unconscious, the skin and diaphragm were opened to expose the heart. A 27G needle (Cat# SV*27EL, Terumo Medical) connected to a 30 mL syringe (Cat# 309650, BD) filled with PBS was inserted into the apex of the left ventricle, and the right ventricle was cut to allow blood to drain. Mice were then perfused with PBS at ∼0.72 mL/min until both testes appeared white (typically 30–60 mL PBS).

For testicular sperm, both testes were collected and decapsulated in EKRB (pH 7.2) containing 1× Krebs-Ringer Bicarbonate Buffer (Cat# K4002, Sigma-Aldrich), 1.26 g/L sodium bicarbonate (Cat# S6761, Sigma-Aldrich), 1× GlutaMAX (Cat# 35050061, Thermo Fisher Scientific), 1× Antibiotic-Antimycotic (Cat# 15240062, Thermo Fisher Scientific), 1× MEM Non-Essential Amino Acids (Cat# 11140050, Thermo Fisher Scientific), and 1× MEM Amino Acids (Cat# 11130051, Thermo Fisher Scientific). After collection, type IV collagenase (Cat# C5138, Sigma-Aldrich) was added to a final concentration of 0.5 mg/mL and samples were incubated at 32°C with rotation for ∼12 min. Intact seminiferous tubules were allowed to settle on ice for ∼5 min, and the supernatant containing testicular sperm was passed through a 100 μm cell strainer, collected, and centrifuged at 800×g for 5 min at 4°C. After one wash in EKRB, the pellet was resuspended and applied to a 2%–4% discontinuous BSA gradient (5 mL of 2% BSA over 5 mL of 4% BSA in a 15 mL tube) and centrifuged at 200×g for 5 min at 4°C. The top 5 mL of supernatant containing purified sperm was carefully collected and centrifuged at 800×g for 5 min at 4°C. Purified sperm pellets were used for downstream experiments.

For caput sperm, both caput epididymides were collected in EKRB. Three incisions were made per caput epididymis, sperm were squeezed out with bent tweezers, passed through a 100 μm cell strainer, centrifuged at 800×g for 5 min at 4°C, and further purified by 2%–4% BSA gradient as described above.

For cauda sperm, both cauda epididymides were collected in 1 mL HTF. Three incisions were made per cauda epididymis, and sperm were allowed to swim up at 37°C for 30 min. Swim-up sperm were collected for downstream separation.

#### Sperm and cytoplasmic droplet (CD) separation

Sperm-containing CDs were subjected to CD purification by discontinuous sucrose gradient centrifugation as previously described^14^ with modifications. Briefly, ∼1 mL of sperm suspension was gently loaded onto a gradient composed of 2 mL of 0.25 M sucrose over 3 mL of 1 M sucrose. After centrifugation at 1,500 ×g for 20 min, the top 2 mL was removed and the subsequent ∼1 mL at the 0.25 M/1 M interface, containing CDs, was transferred to a new tube and centrifuged at 1,200×g for 10 min to pellet residual sperm. The CD-containing supernatant (1 mL) was diluted in 9 mL of 1× PBS and centrifuged at 12,500×g for 20 min at 4°C to enrich CD pellets. The sperm pellet at the bottom of the 1 M sucrose layer was washed once with 1× PBS, treated with somatic cell lysis buffer (SCLB: 0.025% SDS, 0.005% Triton X-100 in 1× PBS), and centrifuged at 3,000×g for 20 min.

#### Surface sensing of translation (SUnSET)

Testicular, caput, and cauda sperm were purified as described above and divided into four treatment groups: (1) no treatment, (2) puromycin only (Cat# A11138-03, Thermo Fisher Scientific), (3) cycloheximide (CHX; Cat# 01810-5G, Sigma-Aldrich, cytoplasmic translation inhibitor) + puromycin, and (4) chloramphenicol (CP; Cat# 227920250, Acros Organics, mitochondrial translation inhibitor) + puromycin. For puromycin treatment, sperm were incubated with 10 μg/mL puromycin at 34°C for 30 min. For CHX + puromycin, sperm were pre-treated with 100 ng/mL CHX at 34°C for 30 min, then pulsed with 10 μg/mL puromycin at 34°C for 30 min. For CP + puromycin, sperm were pre-treated with 25 μg/mL CP at 34°C for 30 min before puromycin addition. Following incubation, samples were washed twice in 1× PBS and processed for immunofluorescence or Western blot.

#### Immunofluorescence

After puromycin incorporation and washing, sperm were spread onto Superfrost Plus slides (Cat# 22-037-246, Thermo Fisher Scientific), air-dried, and fixed in 4% paraformaldehyde (Cat# J19943-K2, Thermo Fisher Scientific) for 15 min. Slides were washed twice in 0.4% Photo-Flo 200/1× PBS (5 min each), once in 0.4% Photo-Flo 200/ddH_2_O (5 min), air-dried, and stored at −80°C. Before staining, slides were equilibrated to room temperature, then immersed in acetone for 20 min at 4°C, rehydrated through graded ethanol (95% twice, 70% twice, 5 min each), and washed three times in 1× PBS. Heat-induced antigen retrieval was performed in citrate buffer (pH 6.0) in a microwave (once at high power for 4 min, three times at low power for 4 min each). After cooling and washing, slides were permeabilized with 0.25% Triton X- 100 (Cat# T8787, Sigma-Aldrich) in 1× PBS for 20 min, then blocked in 1× blocking solution (5% normal donkey serum, 5% fetal bovine serum, 1% BSA in 1× PBS) for 1 h at room temperature. Primary antibodies were applied overnight at 4°C: anti-Puromycin (Cat# MABE343, Sigma-Aldrich, 1:500) and anti-LDHC (Cat# PA5-18779, Thermo Fisher Scientific, 1:100). Secondary antibodies (Alexa Fluor™ Plus 594 donkey anti- mouse, Cat# A32744, and Alexa Fluor™ Plus 488 donkey anti-rabbit, Cat# A32790, both 1:500, Thermo Fisher Scientific) were applied for 1 h at room temperature. Slides were mounted with Antifade Mounting Medium with DAPI (Cat# H-1800, Vector Lab) and imaged on a Nikon ECLIPSE Ti2 confocal microscope with NIS-Elements software.

#### Western blot

Following puromycin incorporation and PBS washes, sperm were lysed in 1× RIPA buffer (Cat# J62725-AP, Thermo Fisher Scientific) supplemented with 1% SDS (Cat# 71736-500ml, Sigma-Aldrich), 357.5 mM 2-mercaptoethanol (Cat# M6250, Sigma- Aldrich), and 1 tablet/10 mL cOmplete™ Mini EDTA-free Protease Inhibitor Cocktail (Cat# 11836170001, Sigma-Aldrich). Lysates were sonicated (3 × 10 s on/20 s off), heated at 95°C for 5 min, and centrifuged at 12,000×g for 20 min. Supernatants were combined with Laemmli buffer (Cat# 1610737, Bio-Rad) containing 2-mercaptoethanol and protease inhibitor, then heated at 100°C for 10 min. Proteins were separated on 4– 20% Mini-PROTEAN® TGX™ Precast Gels (Cat# 4561094, Bio-Rad) and transferred to Amersham™ Protran® nitrocellulose membranes (Cat# GE10600003, Sigma-Aldrich).

Membranes were stained with Ponceau S (Cat# P7170, Sigma-Aldrich) to verify equal loading, de-stained with 0.1 M NaOH, blocked in 5% skim milk in TBST for 1 h, and incubated overnight at 4°C with anti-Puromycin antibody (Cat# MABE343, Sigma- Aldrich, 1:2500 in TBST/5% skim milk). After three TBST washes, HRP-conjugated Rabbit Anti-Mouse IgG (Cat# ab6728, Abcam, 1:5000) was applied for 1 h at room temperature. Bands were detected with the WesternBright ECL kit (Cat# K-12045-D20, Advansta).

#### BONCAT nascent protein enrichment

Purified testicular sperm were divided into homopropargylglycine (HPG)-treated and untreated groups. To deplete intracellular methionine reserves, all samples were cultured at 37°C for 30 min in HPG-free medium (L-methionine-free DMEM [Cat# 21013, Thermo Fisher Scientific] supplemented with 1× GlutaMAX [Cat# 35050061, Thermo Fisher Scientific], 63 mg/L L-cystine dihydrochloride [Cat# RES1513C-A701X, SAFC], and 10% dialyzed fetal bovine serum [Cat# 26400044, Thermo Fisher Scientific]). HPG- group sperm were then incubated with 4 mM L-homopropargylglycine (Cat# 1067-100, Click Chemistry Tools) at 37°C for 2 h; control sperm were cultured in DMEM (Cat# 11995, Thermo Fisher Scientific) with 10% dialyzed FBS for 2 h. After two PBS washes, samples were lysed and sonicated as described above. Clarified supernatants were acetone-precipitated (4:1 ratio, −20°C overnight), pellets were resuspended in 0.1% SDS/PBS with protease inhibitor, and pre-cleared with 50 μL streptavidin beads (Cat# 65601, Thermo Fisher Scientific) for 2 h at room temperature. Endogenously biotinylated proteins retained on beads were eluted in 2% SDS/8 M urea/3 mM biotin in Tris buffer. The HPG-labeled proteins in the supernatant were subjected to a copper-catalyzed click reaction containing 100 μM biotin-azide (Cat# 1265-5, Click Chemistry Tools), 1 mM TCEP (Cat# C4706, Sigma-Aldrich), 100 μM TBTA (Cat# 678937, Sigma-Aldrich), and 1 mM CuSO_₄_ (Cat# 451657, Sigma-Aldrich), acetone-precipitated overnight, and then affinity-purified on streptavidin beads. Beads were washed three times in 0.1% SDS/PBS and twice in 1× PBS. Enriched proteins were eluted in 2% SDS/8 M urea/3 mM biotin in Tris buffer at room temperature for 15 min then 95°C for 15 min. Protein concentration was measured by Bradford assay (Cat# A55866, Thermo Fisher Scientific), and samples were stored at −80°C.

#### SUNsET nascent protein enrichment

Purified CD-less cauda sperm were divided into 5′-azido-puromycin (AP)-treated and untreated groups. The AP group was incubated with 10 μM 5′-azido-puromycin (Cat# CLK-110-S, Jena Bioscience) in EKRB at 34°C for 30 min; control sperm were cultured in EKRB alone at 34°C for 30 min. After two PBS washes, samples were lysed, sonicated, and acetone-precipitated as described above. Pre-cleared supernatants (endogenous biotin removal with streptavidin beads) were reacted overnight with 20 μM DBCO-PEG4-Biotin (Cat# CLK-A105P4-10, Jena Bioscience) in 1× PBS. After acetone precipitation, pellets were resuspended and affinity-purified on streptavidin beads (50 μL/mL sample) as described for BONCAT. Enriched nascent proteins were eluted, quantified by Bradford assay, and stored at −80°C.

#### Proteomics: testicular sperm, caput sperm, cauda sperm, and BONCAT nascent proteins

Protein digestion – Protein content was estimated by bicinchoninic acid assay. 100 μg of protein extract was reduced, alkylated with iodoacetamide, and digested with a trypsin/Lys-C mixture using the EasyPep Mini MS Sample Prep Kit (Cat# A40006, Thermo Fisher Scientific) according to the manufacturer’s protocol. The enzyme-to-protein ratio was 1:10. Peptides were cleaned up using the provided column.

Liquid chromatography – Peptides were analyzed on an UltiMate 3000 RSLCnano system (Thermo Scientific) on a self-packed 100 μm i.d. picofrit column (New Objectives) packed to 30 mm with 1.9 μm ReproSil-Pure C18 (Dr. Maisch). Separation used a 180- min gradient from 2–27% solvent B (solvent A: 0.1% formic acid; solvent B: acetonitrile + 0.1% formic acid) at 50°C with a digital Pico View nanospray source (New Objectives).

Mass spectrometry – Data-independent acquisition (DIA)^60,61^ was performed as described by Searle et al. ^62^ Six gas-phase fractions (GPF1–6, covering 398–1002 m/z) were acquired from a pooled sample to generate a reference spectral library. GPF acquisition used 4 m/z staggered precursor isolation windows at 60,000 resolution, AGC target 4 × 10 , maximum injection time 55 ms, and NCE 33 (HCD). Biological samples were acquired at identical gradient settings using 24 m/z staggered windows over 385– 1015 m/z (60,000 resolution, maximum injection time 55 ms, AGC target 4 × 10 , NCE 33). An empirically corrected library combining GPF and deep neural network Prosit^63^ predictions was used for fragment and retention-time matching. ScaffoldDIA (Proteome Software) or Spectronaut (Biognosys) was used for data processing, requiring exclusive peptide assignment and ≥2 peptides per protein at <1% protein-level FDR. Amica was used for downstream statistical analysis^64^.

#### Proteomics: cauda sperm SUNsET nascent proteins

Protein digestion – Proteins were reduced with dithiothreitol (DTT) at 95°C for 10 min, alkylated with iodoacetamide (IAA) in the dark for 30 min, then precipitated in cold acetone at −20°C overnight. Pellets were collected at 16,000×g for 10 min at 4°C, washed with cold methanol, resuspended in ammonium bicarbonate buffer, and digested overnight at 37°C with trypsin at a 1:30 (enzyme:protein) ratio. Digestion was quenched with formic acid (pH ∼3–4).

LC-MS/MS analysis – Tryptic peptides were analyzed on an Easy-nLC 1200 system (Thermo Fisher Scientific) coupled to an Orbitrap Q Exactive mass spectrometer (Thermo Fisher Scientific) on a PepMap RSLC C18 column (25 cm × 50 μm, 2 μm particle size, 100 Å pore size) at 300 nL/min. A 35-min gradient was used: 0–10% B (0– 2 min), 10–45% B (2–27 min), 45–100% B (27–28 min), 100% B (28–35 min); Buffer A: 0.1% formic acid; Buffer B: 80% acetonitrile + 0.1% formic acid. DIA acquisition parameters: spray voltage 1.7 kV, capillary temperature 300°C; MS1 60,000 resolution, m/z 348–1100, AGC 3 × 10 , injection time 55 ms; MS2 30,000 resolution, AGC 1 × 10 , NCE 30, injection time 55 ms. Variable isolation windows: 25 m/z (m/z 350–400), 20 m/z (m/z 400–870), 40 m/z (m/z 870–1110).

Data processing – Raw data were processed with DIA-NN (v2.1.0; https://github.com/vdemichev/DiaNN) against a UniProt mouse FASTA database. Fixed modifications included carbamidomethylation of cysteine and N-terminal methionine excision; oxidation of methionine was set as a variable modification. Maximum missed cleavages were set to 1. Amica was used for further statistical analysis ^64^.

#### Northern blot

Northern blot was performed as previously described ^65,66^ with modifications. Azide-labeled DNA oligonucleotide probes were ordered from IDT and conjugated to IRDye 680RD DBCO (Cat# 929-50005, Li-Cor) or IRDye 800CW DBCO (Cat# 929-55000, Li-Cor) by incubation in 1× PBS at room temperature in the dark. For small RNAs, 15% TBE-Urea gels (Cat# EC6885BOX, Thermo Fisher Scientific) were pre-run at 300 V for 30 min. 150 ng total RNA was mixed with 2× Formamide Loading Dye (Cat# AM1560, Thermo Fisher Scientific), denatured at 95°C for 3 min, chilled on ice, and separated at 180 V for 1 h in 1× TBE. For large RNAs, 1% agarose gels in 1× NorthernMax™ Denaturing Gel Buffer (Cat# AM8676, Thermo Fisher Scientific) were run at 180 V for 40 min in 1× RNA Gel Buffer (Avantor, Cat# 10128-502). RNA was transferred to BrightStar™-Plus Nylon Membrane (Cat# AM10100, Thermo Fisher Scientific) by Novex™ Semi-Dry Blotter (Cat# SD1000, Thermo Fisher Scientific) in 0.5× TBE at 250 mA for 60 min. Membranes were crosslinked twice at 120 mJ/cm², pre-hybridized in ExpressHyb solution (Cat# 636832, Clontech) at 42°C for 30 min, then hybridized overnight at 42°C with 1 pmol/mL IRDye-labeled probe. After washing in 2× SSC/0.1% SDS and 1× SSC/0.1% SDS, membranes were imaged on a ChemiDoc MP Imaging System (Bio-Rad) and bands were quantified with ImageJ.

#### RNA *in situ* hybridization (sRNA-ISH and RNA FISH)

sRNA-ISH and RNA fluorescence in situ hybridization (FISH) were performed using the miRNAscope™ HD reagent kit (Cat# 324500, ACD Biotechne) and RNAscope Multiplex Fluorescent Assay v2 (Cat# 323110, ACD Biotechne) per the manufacturer’s instructions. Sperm were fixed in 4% paraformaldehyde (Cat# J19943-K2, Thermo Fisher Scientific) for 15 min at room temperature, spread onto Superfrost Plus slides (Cat# 22-037-246, Thermo Fisher Scientific), dried at room temperature, and stored at 4°C. Slides were post-fixed in −20°C-prechilled 100% acetone (Cat# A949-1, Fisher Scientific) for 20 min, washed twice in 100% ethanol (Cat# HC-800-1GAL, Fisher Scientific) for 2 min each with agitation, and post-fixed in 10% normal buffered formalin (Cat# HT501128-4L, Sigma-Aldrich) for 16 h at room temperature. After quenching endogenous peroxidase (RNAscope® Hydrogen Peroxide, 10 min), slides were subjected to target retrieval in 1× RNAscope® Target Retrieval Solution (Cat# 322000, ACD Biotechne) at ≥99°C for 15 min, followed by Protease III or IV treatment at 40°C for 30 min in an ACD HybEZ™ II Hybridization System (Cat# 321710-R, ACD Biotechne).

Hybridization with ACD Biotechne probes was performed at 40°C for 2 h, followed by signal amplification. For sRNA-ISH, red signal was developed, slides were counterstained with 50% hematoxylin (Cat# MHS32-1L, Sigma-Aldrich), blued in 0.02% ammonium hydroxide (Cat# 221228-500ML-PCA, Sigma-Aldrich), dried at 60°C for 15 min, dipped in 100% xylene (Cat# HC-700-1GAL, Fisher Scientific) twice, and mounted with Vectamount (Cat# H-5000, Vector Lab). For large RNA FISH, HRP signal was developed after amplification, and slides were mounted with VECTASHIELD Vibrance® Antifade Mounting Medium with DAPI (Cat# H-1800, Vector Laboratories).

#### Ribo-lite

Purified cells or CHX-treated testicular CDs were lysed in 310 μL of ice-cold lysis buffer containing 20 mM Tris-HCl (pH 7.4), 150 mM NaCl, 5 mM MgCl_₂_, 1 mM DTT, 100 μg/mL CHX, 1% Triton X-100, and 25 U/mL Turbo DNase (Cat# AM2239, Ambion) for 10 min on ice. Lysates were clarified at 20,000×g for 10 min at 4°C, and the supernatant was treated with 1 μL of RNase I (100 U/μL; Cat# AM2295, Ambion) at room temperature for 45 min with gentle rotation. SUPERase•In (10 μL; Cat# AM2696, Ambion) was added, and the sample was overlaid onto 700 μL of 1 M sucrose cushion in lysis buffer (without Triton X-100 and DNase, supplemented with 20 U/mL SUPERase•In). Ribosome-protected fragments (RPFs) were pelleted at 260,000×g for 4 h at 4°C in an MLA-150 rotor (Beckman Optima MAX-XP). The supernatant was discarded, and the pellet was resuspended in 50 μL pellet buffer (10 mM Tris pH 7.5, 1% SDS). RPFs were extracted with TRIzol (Cat# 15596018, Life Technologies) and 200 μL chloroform, precipitated from the aqueous phase with isopropanol and GlycoBlue™ Coprecipitant (Cat# AM9516, Thermo Fisher Scientific) at −20°C overnight, washed in 80% ethanol, and resuspended in 6 μL nuclease-free water.

RPFs (6 μL) were denatured in Gel Loading Buffer II (Cat# AM8546G, Thermo Fisher) at 80°C for 90 s, then separated for 65 min at 200 V on a 15% polyacrylamide TBE-urea gel pre-run at 200 V for 20 min. Samples were loaded in alternating wells to minimize cross-contamination. The region corresponding to 26–34 nt RPFs was excised and confirmed with RNA ladders stained with SYBR™ Gold Nucleic Acid Gel Stain (Cat# S11494, Thermo Fisher Scientific). Gel slices were eluted overnight at room temperature with gentle rotation in 400 μL RNA extraction buffer (300 mM sodium acetate pH 5.5, 1 mM EDTA, 0.25% SDS), snap-frozen at −80°C for 20 min to facilitate extraction, and RNA was recovered by isopropanol precipitation with GlycoBlue™.

Purified RPFs were resuspended in 4 μL nuclease-free water and subjected to library preparation using the D-Plex Small RNA-seq Kit (Cat# C05030001, Diagenode) with D- Plex Unique Dual Indexes (Cat# C05030021, Diagenode). Barcoded libraries (∼203 bp) were size-selected to 175–240 bp using a 3% Agarose Gel Cassette (Cat# BDQ3010, Sage Science) on BluePippin (Cat# PB03352, Sage Science). Final pooled libraries were sequenced on Illumina NovaSeq 6000 with 100 bp single-end reads.

#### Rete testis injection

Injection solutions (1× sterile PBS + 0.02% Trypan Blue [Cat# 15250061, Thermo Fisher Scientific] containing either 0.1% DMSO vehicle or 100 ng/mL CHX + 0.1% DMSO) were prepared fresh on the day of surgery. Adult male recipients were mated with females the day before surgery to clear residual cauda sperm. One rete testis per mouse received 2 μL CHX solution; the contralateral rete testis received 2 μL DMSO vehicle. Only animals with confirmed bilateral injections were included in the analysis. Mice were euthanized 4 days post-injection, and testes and sperm were collected for histological analysis and CASA (SCA Evolution). The paired two-sample t-test was used for the statistical analysis.

#### Cauda sperm chloramphenicol treatment

Cauda epididymides from both sides of each mouse were collected and processed independently. One side was incubated in HTF medium containing 0.1% DMSO vehicle, and the contralateral side was incubated in HTF containing 25 μg/mL chloramphenicol (CP) and 0.1% DMSO. Following incubation at 37°C for 30 min, motility parameters were analyzed by computer-assisted sperm analysis (CASA) using SCA Evolution software. The paired two-sample t-test was used for the statistical analysis.

### QUANTIFICATION AND STATISTICAL ANALYSIS

All quantitative data are expressed as mean ± standard error of the mean (SEM). Statistical significance was determined using Student’s t-test (two groups) or one-way ANOVA (multiple groups) with GraphPad Prism, Microsoft Excel, or R (The R Foundation). Statistical symbols: *, P < 0.05; **, P < 0.01; ***, P < 0.001; ns, not significant. The number of biological replicates for each experiment is indicated in the corresponding figure legends.

## Notes

### Competing Interest Statement

The authors have declared no competing interest.

