## Supplemental Figures and Tables for "Life without cytoplasm: sperm sustain protein production utilizing cytoplasmic droplets and mitochondria as translational apparatuses"

### Supplemental Information

#### Note S1. Supplemental Experimental Procedures: Optimization of Testicular Sperm Isolation (Related to STAR Methods)

Seminiferous tubules are surrounded by collagen, and collagenase digestion releases most, if not all, sperm from the testis tissue. Since sperm typically sediment in ~0.5% BSA gradients during STA-PUT cell sorting[1], we tested STA-PUT cell sorting using discontinuous BSA gradients of 0%–0.5%, 0.5%–1%, 1%–2%, and 2%–4% (Figure S2A). Two hours of gravity sedimentation failed to separate sperm from other cells. We therefore applied a centrifuge force of 200 × g for 5 min. The 0%–0.5% BSA gradient did not separate sperm from other cells, resulting in ~23.3% sperm purity (Figures S2B and S2C). The 0.5%–1% and 1%–2% gradients achieved ~53% sperm purity but retained ~13% and ~14.4% of sperm in the pellet fraction, respectively (Figures S2B and S2C). The 2%–4% discontinuous BSA gradient with 200 × g centrifugation showed the lowest pellet retention (~10% sperm) and a purity comparable to other gradients (~51%) (Figure S2B and S2C). Because the primary contaminants were red blood cells (Figure S2B), we introduced cardiac perfusion with PBS prior to testis collection to remove erythrocytes. Following perfusion, the top 5 mL of supernatant from the 2%–4% discontinuous BSA gradient (200 × g, 5 min) yielded ~70% sperm purity (Figure S2E and S2F). A 4%–8% gradient further improved purity to ~80%, but at the cost of CD loss (Figure S2F). We therefore adopted the final protocol: cardiac perfusion → 2%–4% discontinuous BSA gradient at 200 × g for 5 min → collection of the top 5 mL supernatant. The same procedure was applied to caput sperm following mechanical extrusion from the caput epididymis (Figure S2F). This protocol also improved cauda sperm purity from ~91.4% to ~98.2% (Figure S2F). Final purities: testicular sperm ~69.3%, caput sperm ~70.4%, cauda sperm ~98.2% (Figure S2G).

**Figure S1. Optimization of Sperm Protein Extraction (Related to STAR Methods)**

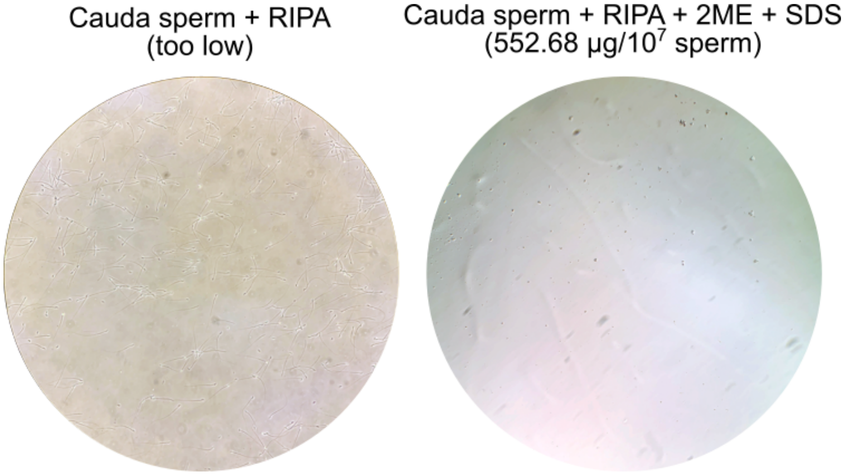


Sperm lysis efficiency with or without supplementation of 2-mercaptoethanol (2-ME) and SDS, assessed by total protein yield. Protein amounts are indicated in parentheses.

### Figure S2. Optimization of Sperm Purification (Related to STAR Methods)

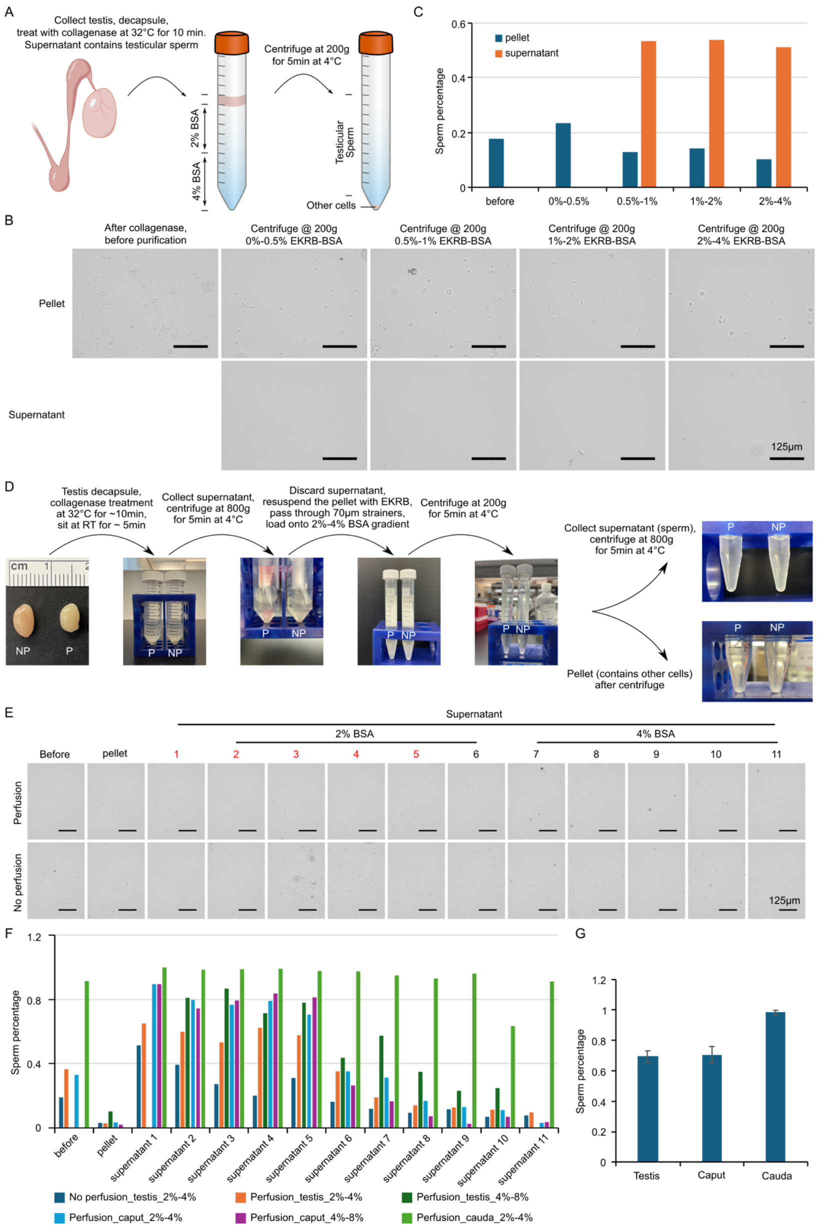

**(A)** Overview of the sperm purification workflow.

**(B)** Representative images of sperm purification fractions at different BSA gradient concentrations.

**(C)** Summary of sperm purity (%) achieved at each BSA gradient concentration.

**(D)** Overview of the sperm purification workflow with and without cardiac perfusion.

**(E)** Representative images of sperm purification fractions with and without perfusion at different BSA gradients.

**(F)** Summary of sperm purity (%) at different supernatant layers from BSA gradients, with and without perfusion.

**(G)** Summary of overall testicular, caput, and cauda sperm purity achieved with the 2%–4% BSA gradient protocol.

**Figure S3. SUnSET in Isolated Cytoplasmic Droplets and CD-Less Sperm (Related to Figure 2)**

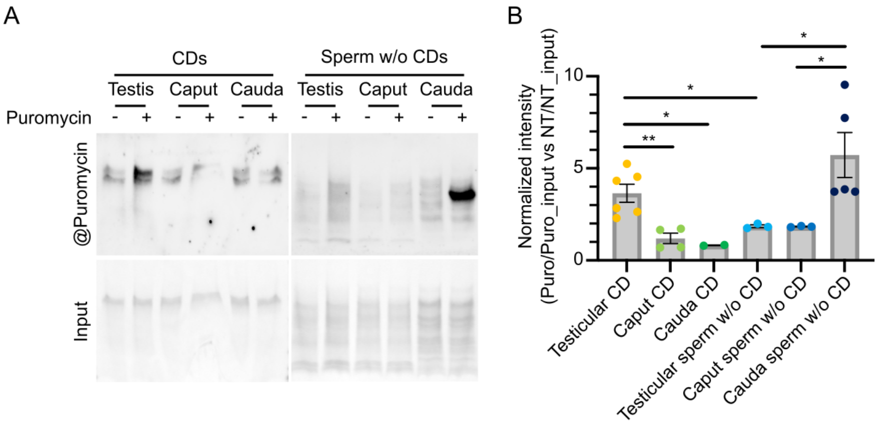


CHX, cycloheximide; CP, chloramphenicol.

**(A)** Representative western blot showing puromycin incorporation efficiency in isolated CDs and CD-less sperm from testicular, caput, and cauda epididymis, with and without inhibitor pre-treatment.

**(B)** Densitometric quantification of western blot signals. Values were normalized to input loading, then the puromycin-treated group was normalized to the corresponding no-treatment control.

**Figure S4. RNA Profiles and Ribo-lite Ribosome-Protected Fragments in Testicular CDs, Caput CDs, Cauda CDs, Testicular Pellets, and Caput Pellets (Related to Figure 4)**

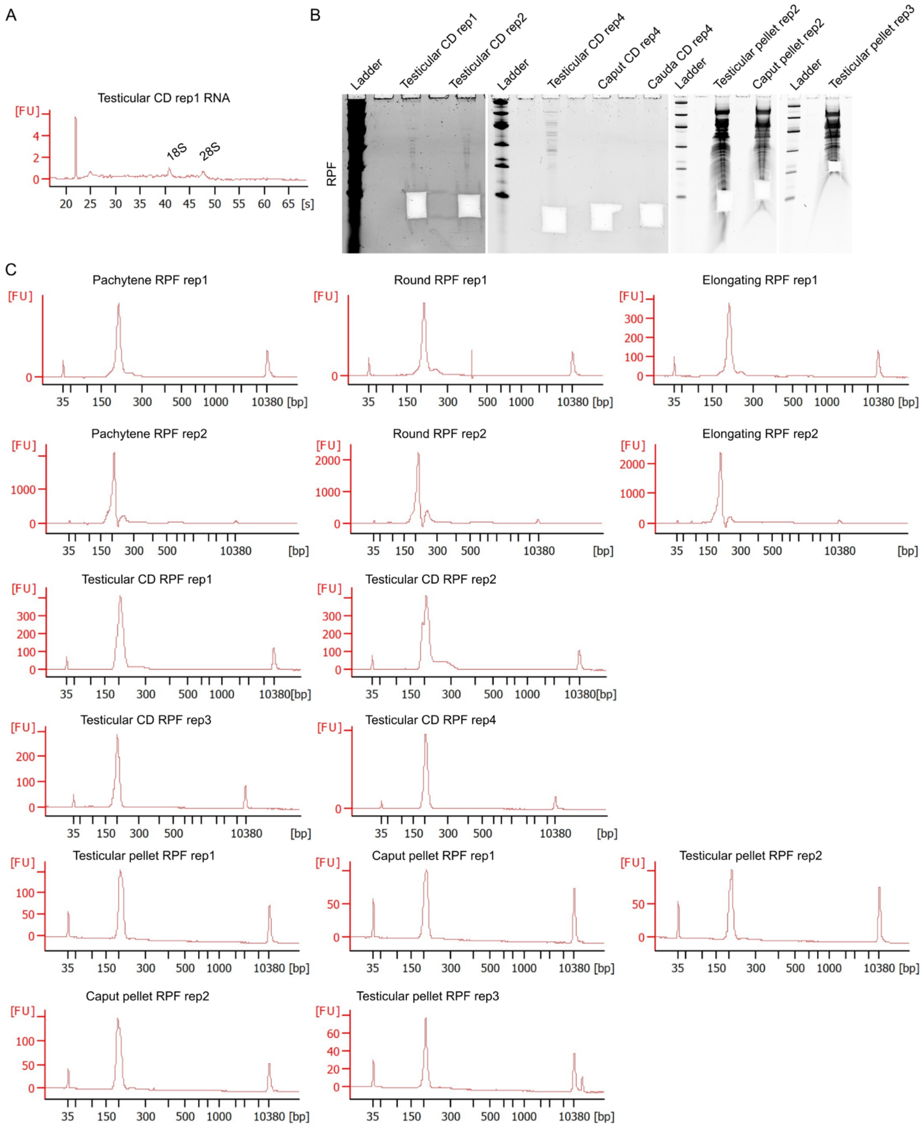


**(A)** Agilent Bioanalyzer RNA profiles of testicular CDs showing the presence of 28S and 18S rRNAs.

**(B)** Gel images of Ribo-lite ribosome-protected fragments (RPFs) after RNase I digestion of testicular CDs, caput CDs, cauda CDs, testicular pellets, and caput pellets.

**(C)** Agilent Bioanalyzer profiles of Ribo-lite libraries prepared from pachytene spermatocytes, round spermatids, elongating/elongated spermatids, testicular CDs, caput CDs, cauda CDs, testicular pellets, and caput pellets, confirming appropriate library size distribution.

**Figure S5. Proteomics of De Novo Synthesized Proteins in Testicular CDs, Testicular Sperm, and Cauda Sperm (Related to Figures 4 and 6)**

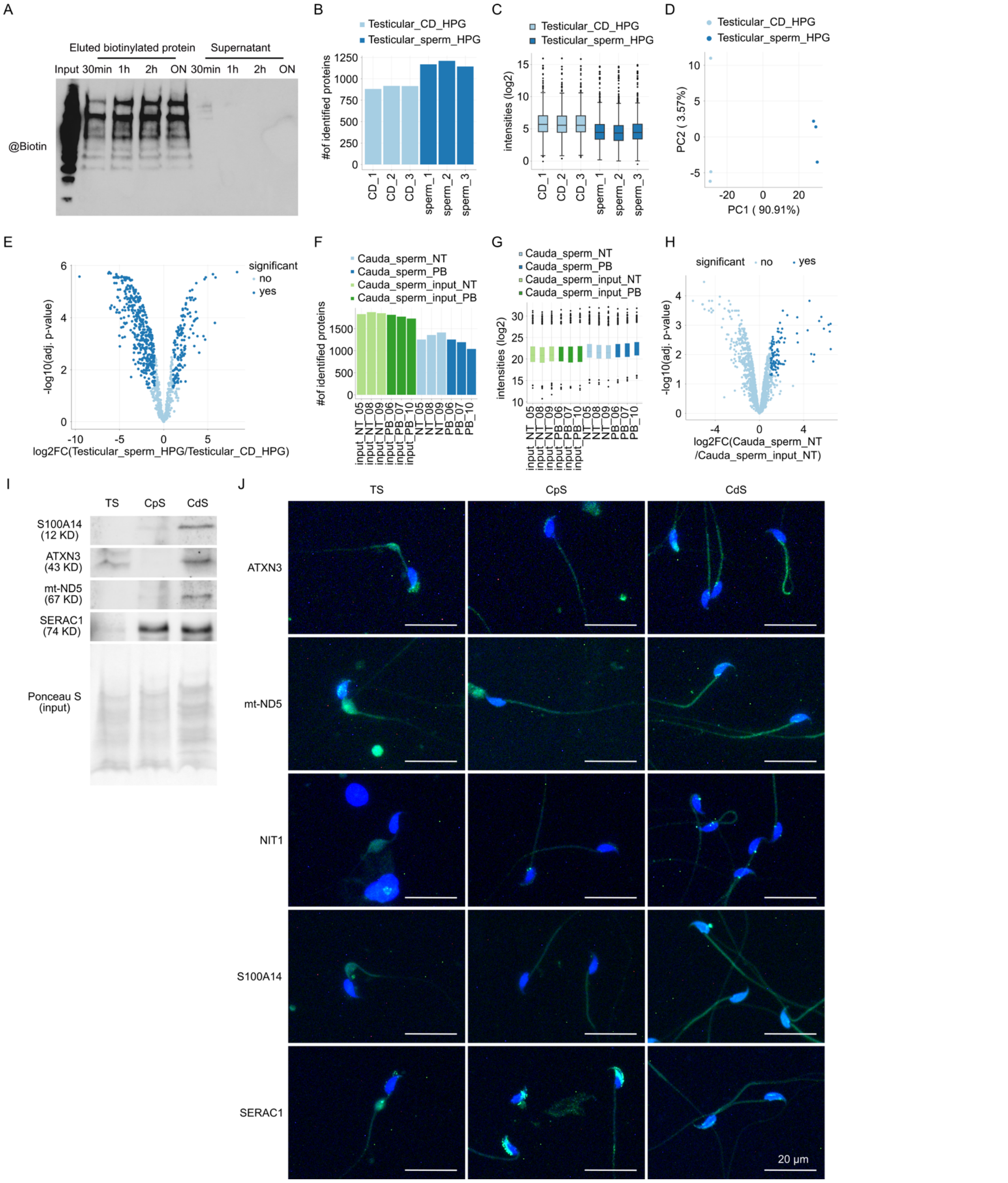


**(A)** Optimization of streptavidin bead pull-down efficiency for biotinylated proteins. Signal intensity as a function of incubation time; 2 h was selected as the optimal condition.

**(B)** Number of BONCAT-identified proteins in testicular CDs and testicular sperm, with and without HPG labeling.

**(C)** Log2 intensities of BONCAT-identified proteins in testicular CDs and testicular sperm.

**(D)** Principal component analysis (PCA) of nascent proteomes from testicular CDs and testicular sperm.

**(E)** Volcano plot showing differentially enriched nascent proteins in testicular CDs versus testicular sperm.

**(F)** Number of SUnSET-identified proteins in cauda sperm treated with puromycin-biotin (PB) or no treatment (NT).

**(G)** Log2 intensities of SUnSET-identified proteins in PB-treated and NT cauda sperm.

**(H)** Volcano plot showing NT-enriched proteins relative to NT-input in cauda sperm.

**(I)** Western blot validation of selected PB-enriched proteins in testicular sperm (TS), caput sperm (CpS), and cauda sperm (CdS). Scale bar = 20 μm.

**(J)** Immunofluorescence localization of selected PB-enriched proteins in TS, CpS, and CdS. Scale bar = 20 μm.

**Supplemental Tables (provided as separate Excel files)**

**Table S1. Proteomes of testicular, caput, and cauda epididymal sperm (Related to Figure 1).** LC-MS/MS-based proteomics of highly purified testicular sperm (3,597 proteins), caput epididymal sperm (2,750 proteins), and cauda epididymal sperm (2,391 proteins), together with the lists of up- and down-regulated proteins identified in the testis→caput and caput→cauda comparisons (log2 fold change ≥ 2).

**Table S2. GO term enrichment of dysregulated sperm proteins during epididymal transit (Related to Figure 1).** GO biological process enrichment for (S2A) proteins upregulated in caput versus testicular sperm, (S2B) proteins downregulated in caput versus testicular sperm, (S2C) proteins upregulated in cauda versus caput sperm, and (S2D) proteins downregulated in cauda versus caput sperm.

**Table S3. Cytoplasmic droplet (CD) proteome (Related to Figures 1 and 3).** LC-MS/MS-based proteomics of purified testicular CDs (1,833 proteins), including the subset of 117 proteins annotated to the cytoplasmic translational machinery (60S and 40S ribosomal proteins, translation initiation factors, aminoacyl-tRNA ligases, elongation factors, and others).

**Table S4. Ribo-lite ribosome-protected fragment (RPF) quantification (Related to Figure 4).** RPKM values of Ribo-lite libraries prepared from pachytene spermatocytes, round spermatids, elongating/elongated spermatids, testicular CDs, caput CDs, cauda CDs, testicular pellets, and caput pellets.

**Table S5. GO term enrichment of Ribo-lite translatomes (Related to Figure 4).** GO biological process enrichment for Ribo-lite translatomes of (S5A) pachytene spermatocytes, (S5B) round spermatids, (S5C) elongating/elongated spermatids, (S5D) testicular pellets, (S5E) genes commonly detected in pachytene spermatocytes, round spermatids, and testicular pellets, (S5F) testicular CDs, and (S5G) caput pellets.

**Table S6. BONCAT-identified nascent proteins in testicular CDs and testicular sperm (Related to Figure 4).** Label-free quantification of proteins enriched by streptavidin pull-down following HPG labeling and click chemistry, together with no-HPG control samples, used to define the testicular CD and sperm nascent proteomes.

**Table S7. GO term enrichment of BONCAT nascent proteomes (Related to Figure 4).** GO biological process enrichment for (S7A) nascent proteins enriched in testicular CDs, (S7B) nascent proteins enriched in testicular sperm, and (S7C) genes shared between the BONCAT nascent proteomes and the Ribo-lite testicular CD translatome.

**Table S8. Nascent proteins identified in cauda epididymal sperm by puromycin-biotin SUnSET proteomics (Related to Figure 6).** Label-free quantification of proteins enriched from cauda sperm incubated with 5′-azido-puromycin (PB) or no treatment (NT), together with the corresponding input samples, used to define the cauda sperm nascent proteome.
